# Chronic high glyphosate exposure delays individual worker bee (*Apis mellifera* L.) development under field conditions

**DOI:** 10.1101/2020.07.08.194530

**Authors:** Richard Odemer, Abdulrahim T. Alkassab, Gabriela Bischoff, Malte Frommberger, Anna Wernecke, Ina P. Wirtz, Jens Pistorius, Franziska Odemer

## Abstract

The ongoing debate about glyphosate-based herbicides (GBH) and their implications for beneficial arthropods give rise to controversy. This research was carried out to cover possible sublethal GBH effects on brood and colony development, adult survival, and overwintering success of honey bees (*Apis mellifera* L.) under field conditions. Residues in bee relevant matrices such as nectar, pollen and plants were measured in addition. To address these questions, we adopted four independent study approaches. For brood effects and survival, we orally exposed mini-hives housed in the “Kieler mating-nuc” system to sublethal concentrations of 4.8 mg glyphosate/kg (T1, low) and 137.6 mg glyphosate/kg (T2, high) over the period of one brood cycle (21 days). Brood development and colony conditions were assessed after a modified OECD method (No. 75). For adult survival, we weighed and labeled freshly emerged workers from exposed colonies and introduced them into non-contaminated mini-hives to monitor life span for 25 consecutive days. Results from these experiments showed a trivial effect of GBH on colony conditions and survival of individual workers, even though hatching weight was reduced in T2. The brood termination rate (BTR) in the T2 treatment, however, was more than doubled (49.84%) when compared to the control (22.11%) or T1 (20.69%). This was surprising as T2 colonies gained similar weight and similar numbers of bees per colony compared to the control, indicating equal performance. Obviously, the brood development in T2 was not “terminated” as expected by the OECD method terminology but rather “slowed down” for an unknown period of time. In light of these findings, we suggest that chronic high GBH exposure is capable of delaying worker brood development to a significant extent while no further detrimental effects seem to appear at the colony level. Against this background, we discuss additional results and possible consequences of GBH for honey bee health.

## 1 INTRODUCTION

With its first commercial glyphosate-based formulation, the Monsanto Chemical Company introduced Roundup® to the market in 1974. This non-selective herbicide was mainly used as weed killer around power lines, train tracks and in specific farming operations. In 1996, Roundup Ready® seeds became available to farmers and worldwide glyphosate use substantially increased. Monsanto’s glyphosate patent expired in 2002, leading to a drop in prices due to growing competition on the market. This development, its high efficiency, and the easy-to-apply practice have led to the increasing use of glyphosate-based herbicides (GBH) during the past three decades (reviewed in Benbrook 2016 and Richmond 2018). Agricultural utilization of GBH includes but is not limited to pre-sowing, pre-harvest (e.g. desiccation) and stubble application. From a farmer’s perspective, therefore, glyphosate primarily reduces labor and machine costs whereas other pesticides intend to improve crop yields. Accordingly, GBH are far more than just a weed killer and must be seen as an agronomical tool (Steinmann et al. 2012).

GBH are usually not applied during flowering by reason of above-mentioned utilization, and yet field crops represent a potential source for honey bee exposure. Due to the control of weeds within or adjacent to these fields, pollinators can become unintentional targets of control measures. Further, Dickeduisberg et al. (2012) provide evidence that grassland conversion could also be a possible source of bees’ exposure. To date, however, there are only few studies reporting field relevant GBH residues in bee products. In Hawaii for example, 27% of analyzed local honey was tested positive (n=59) with an average of 118 mg/kg glyphosate per sample (Berg et al. 2018). In Europe, this honey would have to be withdrawn from the market as the maximum residue level (MRL) of glyphosate is limited to 0.05 mg/kg (EC, 2020). A similar range of contamination was found in Estonia in 2013 (22% of analyzed honeys, n=33), however to a many-fold lower degree (0.044 mg/kg) (Karise et al. 2017). In a worldwide survey, 59% from a total of 69 honey samples contained glyphosate residues ranging from 0.017 to 0.163 mg/kg (Rubio et al. 2014). Even though these are important findings, data are not sufficient to derive a realistic exposure scenario for honey bees. Metabolites, for instance, are most often not adequately considered. They may produce unexpected effects or potential toxicities and, therefore, it is important to identify them (Stevens & Sumner 1982). Hence, the major bacterial metabolite of glyphosate aminomethylphosphonic acid (AMPA) deserves particular attention in such a scenario.

Because glyphosate’s target enzyme 5-enolpyruvylshikimate 3-phosphate synthase (EPSPS) is not found in insects or other animals, it is generally considered not harmful to these organisms (Schönbrunn et al. 2001). In accordance with results from regulatory testing, a risk for honey bees could not be identified and, therefore, GBH are classified as “not harmful to bees” (B4) in Germany. Common endpoints of ecotoxicological studies in the laboratory usually relate to acute toxicity within a certain period of time. They are indicated either by the lethal dose (LD_50_) or lethal concentration (LC_50_) that causes death in 50 percent of the treated individuals. Thus, high toxicity results in low LD_50_/LC_50_ values and *vice versa*. For honey bees, the oral and contact LD_50_ for glyphosate are determined to be 100 and >100 µg/bee suggesting a low acute toxicity (EFSA, 2015). More recently, however, several studies conducted under laboratory conditions reported sublethal effects of the herbicide. Besides impaired cognitive abilities and navigation skills (Balbuena et al. 2015), primarily implications for the host-microbiome were indicated (Motta et al. 2018, Dai et al. 2018, Blot et al. 2019). While microbiomes’ consequences for bee health are still under discussion, infliction of harm to physiological parameters give more rise to concern at this stage. In worker bees, GBH seem to hypertrophy and damage not only royal jelly-producing glands (Faita et al. 2018), but also delay larval molting and reduce larval weight suggesting serious implications at the colony level (Dai et al. 2018, Vazquez et al. 2018). Prolonged brood development could, for example, entail a higher reproductive success of *Varroa destructor*, to name just one (Frey et al. 2013, Odemer 2020a).

In a previous study, Thompson et al. (2014) set up a large field trial to test GBH at the colony level and in particular on brood development. Field residue data were used to create a realistic worst-case scenario for acute exposure. Free-flying colonies were fed 1L sucrose solution treated with technical grade glyphosate. Brood development was assessed until day 16 without evidence for detrimental effects, neither on brood and adult mortality nor on pupal weight. However, a full brood cycle or beyond - i.e. adult survival after chronic exposure - was yet not covered. Hence, the impact of GBH on colony performance parameters such as brood development or overwintering success under chronic conditions still needs to be answered. This study seeks to obtain data, which will help to address these research gaps. In different approaches, we thus examined a broad array of performance parameters at the combined individual and colony level. According to current knowledge, we hypothesized that (1) larval weight loss should be translated to adult workers after chronic exposure, (2) if survival and/or overwintering are affected, clearly visible signs should emerge by close examination of colony conditions and (3) delayed molting must be reflected by brood development to some degree. In addition, we wanted to confirm/expand (4) field-realistic GBH residue data in different bee matrices to complete the picture of a realistic exposure scenario.

## 2 MATERIALS & METHODS

This research was carried out including four different study approaches:

(2.1) Experiment 1: *Brood development and survival*
(2.2) Experiment 2: *Field exposure*
(2.3) Experiment 3: *Overwintering*
(2.4) Experiment 4: *Field residues*

### 2.1 Brood development and survival

This field study was conducted from June to September 2019.

#### 2.1.1 Experimental colonies and field site

Here we used the “Kieler mating-nuc” system, a Styrofoam box with four top-bars, a strip of a beeswax foundation attached to it and equipped with a feeder (Fig. 1), recently presented in Odemer et al. (2018). In brief, 19 mini-hives were established with about 800 worker bees originating from brood frames of two healthy donor colonies with no clinical symptoms of adult bee or brood diseases visible during inspection. Subsequently, unmated sister queens (*Apis mellifera* L.) were introduced and the mini-hives were placed at a remote apiary near Wurmberg, Germany for mating after confinement in a dark and chilled room for 24 h. Corresponding coordinates were as follows: latitude 48.867414°, longitude 8.808538°. Within a period of five weeks, the established hives showed all relevant brood stages (eggs, larvae, and patches of sealed brood) and newly built combs as a sign of successful mating. In total, fifteen of the mini-hives were randomly assigned to the control and the two treatments T1 and T2, five (= replicates) to each group, respectively. One T2 mini-hive was discovered queenless shortly after the start of the experiment and was, therefore, removed (n: C=5, T1=5, T2=4). We maintained the remaining four mini-hives from the total of 19 for the survival assessment of the experiment (see 2.1.5).

**Fig. 1.**
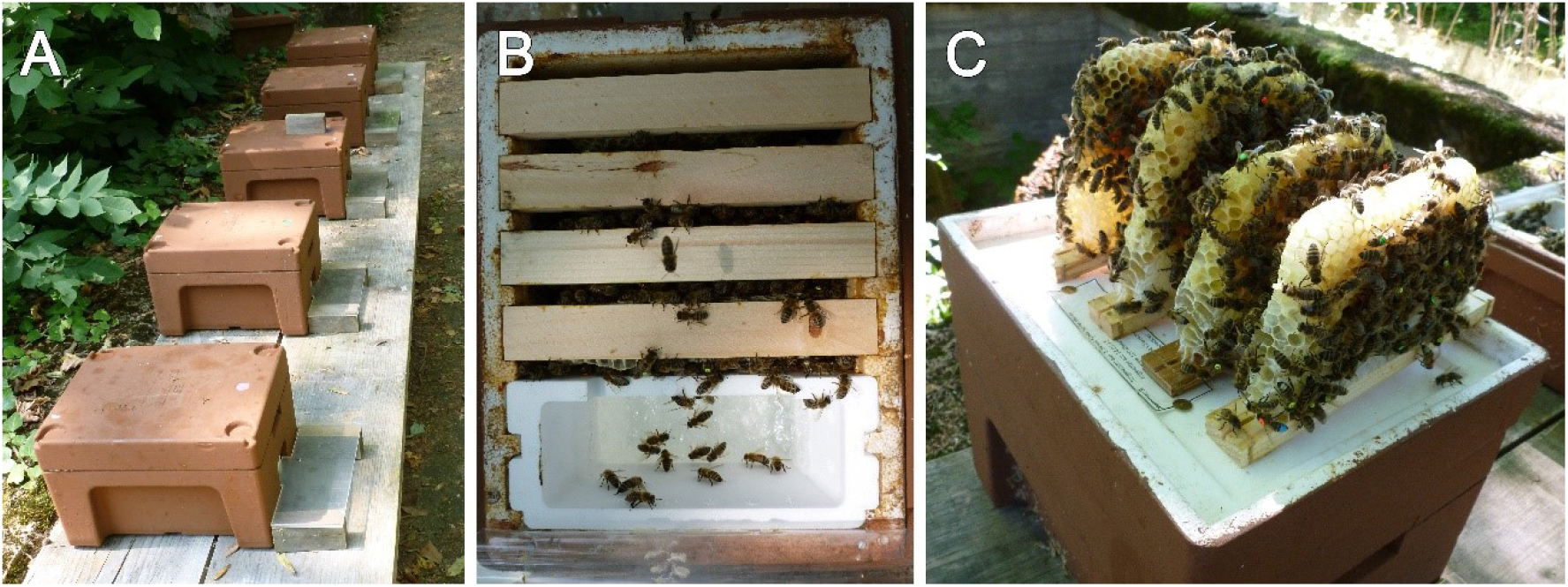
**A** Mini-hives of one treatment group including five colonies during glyphosate exposure. **B:** plan view of a mini-hive with four top-bars and a feeding device, providing space for an intact honey bee colony. **C:** for the brood/mortality assessment, all four combs were removed and both sides of each top-bar without/with bees attached were photographed, respectively (see 2.1.4 and 2.1.5)]

Within the proximity of 250 m, no other hives were set up whereas in an extended radius (> 250 m) a sufficient number of unrelated colonies were located to provide enough drones for mating. At the time present, bees could forage on *Tilia* spp*., Cyanus segetum* and other floral sources in the surroundings as the weather provided favorable conditions (avg. outdoor temperature 18.9 °C, precipitation 53.97 L/m^2^, Wetter BW 2019).

#### 2.1.2 Chemical treatment and sampling

Glyfos Unkraut-Frei® (Dr. Stähler, Germany) with a glyphosate concentration of 360 g a.i./L was used for the treatment. Accordingly, the product was directly mixed into 5 L of feeding syrup (Apiinvert, Südzucker GmbH, Germany) to achieve the desired concentrations. The treatment was comprised of two feeding regimes; T1 with a measured concentration of 4.8 mg a.i./kg and T2 with a measured concentration of 137.6 mg a.i./kg feeding syrup. This equals to 0.090 g product/5L for T1 and 2.610 g product/5L for T2. The control was fed with untreated syrup like all other groups in a weekly interval. Chronic treatment in T1 and T2 was maintained for 21 consecutive days with a total amount of 0.81 kg feeding syrup per hive. This corresponds to a calculated amount of T1: 3.859 mg and T2: 111.428 mg glyphosate per hive, respectively.

Ahead of the experiment, a sample of each feeding regime and the untreated feeding syrup was collected, respectively. On day 21, pooled samples of stored syrup from all groups were collected from in-hive storage cells for residue analysis.

#### 2.1.3 Colony conditions

The total number of bees was estimated for each colony according to the “Liebefeld method” (Imdorf et al. 1987). In addition, the absolute hive weight was recorded at the time of the colony assessments with a precision scale (Mettler-Toledo Kompaktwaage ICS445k, Germany, resolution 0.01 g). The assessments were performed in July (31^st^), August (30^th^) and September (25^th^).

#### 2.1.4 Brood development and photographic assessment

At the colony condition assessment before the first application of the test item, from groups C, T1, and T2 (Brood Area Fixing Day = BFD), one or several brood combs were taken out of each replicate. The development of the bee brood was continuously assessed and followed by selecting 100 cells per replicate containing eggs, approximately. Pictures were taken from the comb-area containing eggs, modified according to Schur et al. (2003). In brief, selected combs were uniquely identified and the fixed brood area was photographed during each brood stage assessment (BFD assessments, Canon EOS 700D with Canon EF-S 24mm F2.8 pancake lens). By this means, each selected cell was identified to evaluate its content on the digital images to follow the process of development. Cells were classified and evaluated following the scheme in Tab. S1. From this evaluation, the brood termination rate (BTR) was calculated, respectively. Further was the original method modified, so that only three assessments were necessary to reduce invasive operations and disturbance of the colonies.

#### 2.1.5 Survival of individual bees and hatching weight

After the 21 d chronic exposure, one sealed brood comb containing brood cells ready to hatch was removed from mini-hives of all groups and put together treatment wise - five combs from C, T1 and four from T2 - in an incubator for 24 hours (Memmert IN30, Germany at 33 °C, 70% RH, total darkness). One replicate from T2 was found to be queenless during the exposure phase and was therefore excluded from further analysis. Subsequently, all hatched bees were pooled treatment-wise and 30-40 young workers were collected at random. For identification, a colored and numbered opalith plate was glued on the thorax using shellac. In addition, we marked the dorsal side of the abdomen with a hive specific color (Posca Marking Pen, Japan) in order to identify drifting bees that enter neighboring colonies. The bees were then introduced into four of the mini-hives (= replicates) remaining from the initial 19, divided equally along their affiliation. The total number (n) of introduced individuals per treatment was as follows: C (152), T1 (149) and T2 (141). Then, survival was monitored for a period of 25 days. See Fig. S2 for a detailed experimental setup scheme. The monitoring started 24 hours after the bees’ introduction. The mortality assessment included a check every two to three days, for which all combs including the inside of the hive were photographed for the later on counting of the marked bees on a computer screen. The pictures were taken outside the foraging activity, early in the morning.

Before labeling of the above-mentioned worker bees, their hatching weight was documented at random with a portable precision scale (Kern CM 60-2N, Germany, resolution 0.01 g). Therefore, single bees were transferred carefully into a in a tared container. They were weighed according to their treatment and respective pool resulting in a total number (n) of C (50), T1 (38) and T2 (36).

### 2.2 Field exposure

This field study was conducted from July to August 2016.

#### 2.2.1 Experimental colonies and field sites

Twelve honey bee colonies (*A. mellifera*), healthy and queen-right with one hive body including ten combs were used. The colonies were as homogeneous as possible at a strength of approximately 15,000 bees per colony. Queens originated from one breeding line (sisters, reared at the test facility in the same year). No clinical symptoms of adult bee or brood diseases were visible during inspection. Colonies were split into two groups with six replicates each (control C and treatment T).

Eight days before application (Days After Treatment = DAT-8) the colonies were placed at two flowering *Phacelia tanacetifolia* plots near the city of Braunschweig with an area of approximately 1 ha, respectively. Corresponding coordinates were as follows: Control plot C; latitude 52.296415°, longitude 10.437062°, Treatment plot T; latitude 52.202098°, longitude 10.623331°. Colonies from the treatment plot were migrated after nine days of exposure (DAT+8) to the control plot C.

#### 2.2.2 Chemical treatment and sampling

Roundup®PowerFlex (Monsanto Agrar Deutschland GmbH) with a glyphosate concentration of 480 g a.i./L was used for the treatment. The treatment was applied with a rate of 3.75 L/ha in 300 L water/ha on the flowering phacelia (BBCH 64-65) plot T (DAT0) with a field sprayer. The control remained unsprayed.

A total of 12 beebread, 60 food, and 10 plant samples (whole plant except the roots) were taken at different time intervals before and after the treatment from both plot sites for residue analysis.

#### 2.2.3 Colony conditions

On three dates, colony conditions were assessed using the “Liebefeld method” (Imdorf et al. 1987). On DAT-6 before the exposure and on DAT+15 and DAT+57 after the exposure, respectively. The time intervals were chosen to cover colony conditions before and after the treatment and to include more than one brood cycle. Parameters such as number of bees, brood cells and stores (honey and pollen) were estimated according to the method.

### 2.3 Overwintering

This field study was conducted from October 2018 until March of the following year.

#### 2.3.1 Experimental colonies and field sites

The same setup as in 2.2.1 was applied, with one exception. Colonies were at a strength of approximately 15,000 bees per colony.

All colonies were located at a remote apiary near the city of Braunschweig. Corresponding coordinates were as follows: latitude 52.202098°, longitude 10.623331°.

#### 2.3.2 Chemical treatment and sampling

Roundup®PowerFlex (Monsanto Agrar Deutschland GmbH) with a glyphosate concentration of 480 g a.i./L was used for the treatment. The treatment was comprised of one feeding regime; T with a nominal concentration of 8.13 mg a.i./kg (according to Motta et al. 2018). Accordingly, the product was directly mixed into 5 L of the feeding syrup (1:1 sugar water, w/w, density: 1.2296) for each colony to achieve the desired concentration, respectively. The control was fed with untreated syrup. Chronic treatment in T was maintained until the feeder was emptied. The average measured concentration of 5.439 mg a.i./kg in the feeding solution corresponds to a calculated total amount of T: 33.436 mg glyphosate per hive.

A total of 66 samples of stored food from all groups were collected from in-hive storage cells for residue analysis at different time intervals.

#### 2.3.3 Colony conditions

On three dates, colony conditions were assessed using the “Liebefeld method” (Imdorf et al. 1987). On DAT-1 before the exposure and on DAT+43 and DAT+170 after the exposure, respectively. The time intervals were chosen to cover colony conditions before and after the treatment, with a focus on during and after overwintering. Parameters such as number of bees, number of brood cells and the number of stores were estimated according to the method.

### 2.4 Determination of GBH residues under semi-field conditions

This semi-field study was conducted from July to August 2017.

#### 2.4.1 Experimental colonies and field sites

The same setup as in 2.2.1 was applied, with the following exceptions. Queens were removed on DAT-14 to stimulate foraging in the tunnel tents. Colonies were split into two groups with three replicates each (control C and treatment T).

On the day of application (DAT0), each colony was placed in a tunnel tent with an area of approximately 33.5 m^2^ on a flowering *P. tanacetifolia* plot at our field site in Braunschweig, respectively. Corresponding coordinates were as follows: latitude 52.296415°, longitude 10.437062°. Due to loss of forage in the GBH treated tents, all colonies were migrated after 14 days of exposure (DAT+13) to a remote apiary.

#### 2.4.2 Chemical treatment and sampling

Roundup®PowerFlex (Monsanto Agrar Deutschland GmbH) with a glyphosate concentration of 480 g a.i./L was used for the treatment. The treatment in the T tents was applied with a rate of 3.75 L/ha in 300 L water/ha (DAT0) on the flowering phacelia (BBCH 65) using a portable boom sprayer (Schachtner, Germany). The control was sprayed with water.

A total of four stored food, two pooled honey sac, two pooled corbicular pollen, and four pooled plant samples were taken at different time intervals after the treatment from all tunnels for residue analysis. On DAT+5, plants could no longer be sampled due to the action of the herbicide.

### 2.5 Residue analysis

The samples were analyzed using a method without derivatization step based on the methods for honey, milk, soybeans and maize published by Chamkasem et al. (2015, 2016, 2016, 2017). Depending on the sample material, some modifications were necessary. The method was validated by analyzing a series of spiked replicate honey, pollen and plants (Phacelia) samples (see supplementary Method S2).

LC-MS/MS was used to identify and quantify glyphosate (GLY) and its metabolite aminomethylphosphonic acid (AMPA) in the samples. The system used was a Nexera X2 HPLC system (SHIMADZU Corp., Kyoto, Japan) coupled to a triple quadrupole mass spectrometer Q TRAP 6500+ (SCIEX, Framingham, MA, USA) equipped with an electrospray ionization (ESI) source. The mass spectrometer was operated in the negative ESI mode and three multiple reaction monitoring (MRM) transitions were monitored for each analyte in order to confirm compound identity.

In undiluted and 1:10 diluted samples, the analyte contents were determined using matrix-matched standards. If samples had to be diluted 1:100 or 1:1000, the analytes were quantified using reference standards in extracting agent as matrix effects were sufficiently reduced by dilution.

For quantification, the internal standard method was used, with Glyphosate-^13^C2^15^N and AMPA^13^C^15^N as internal standards (SANTE, 2019). This procedure minimizes the matrix effect and enables accurate quantification.

The average recoveries in plants and pollen at the fortification levels of 25, 50 and 250 μg/kg were between 73% and 83% for GLY and 71% and 87% for AMPA with relative standard deviations (RSD) of less than 20% for both analytes. An exception was the recovery of 148% (RSD 18%) for GLY in pollen at the limit of quantification of 25 μg/kg.

For GLY and AMPA the average recoveries in honey at the fortification levels of 25 μg/kg, 250 μg/kg and 50 mg/kg were in the range of 74% and 112% with RSD between 9% and 17%.

In honey and plants the LOD of GLY was 5.0 μg/kg and in pollen 12.5 μg/kg, respectively. The LOQ was 12.5 μg/kg and 25 μg/kg, respectively.

In honey the LOD of AMPA was 2.5 μg/kg and in plants and pollen 12.5 μg/kg, respectively. The LOQ were 5.0 μg/kg and 25 μg/kg, respectively.

For details of the analytical method and the method validation, see Method S2 in the supplementary material.

### 2.6 Statistical analysis

The estimated number of bees and brood cells from all colony condition assessments, hatching weight and weight gain of the mini-hives as well as brood termination rate, and the number of selected eggs from the BFD were checked with a Shapiro-Wilk test for normal distribution. If data were normal distributed, either a student’s t-test or a one-way ANOVA was performed to compare experimental groups, respectively. Where pairwise tests were performed, *P*-values were corrected with the Bonferroni method. Mortality in the mini-hives was evaluated with a Kaplan-Meier-Survival analysis (KM). Survivorship between control and treatments was compared pairwise and tested for significance with Log-Rank tests (Cox-Mantel). Individuals remaining alive at the end of the experiment were considered censored, as were those observed but not collected on the final day. All groups that underwent the KM analysis including the four replicates used in the monitoring phase were additionally compared using a Cox proportional hazards model to determine the hazard ratio (HR). Possible inter-colony effects were evaluated as a covariate to justify data pooling of the same treatments. For all tests, RStudio (R Core Team, 2019) and a significance level of α= 0.05 was used.

## 3 RESULTS

### 3.1 Brood development and survival

#### 3.1.1 Colony conditions

Because of the complete randomized assignment of the mini-hives to their respective treatment, initial colony strength differed slightly but not significantly between groups with control C being the weakest (Fig. 3A, Jul, *P* > 0.05, ANOVA). However, group C demonstrated compensation in the following two assessments leveling out these differences in the median number of bees (Fig. 3A, Jul, Aug). With the progression of the study, absolute colony weight was similarly increasing with no significant differences between groups (Fig. 3B, *P* > 0.05, ANOVA).

**Fig. 2.**
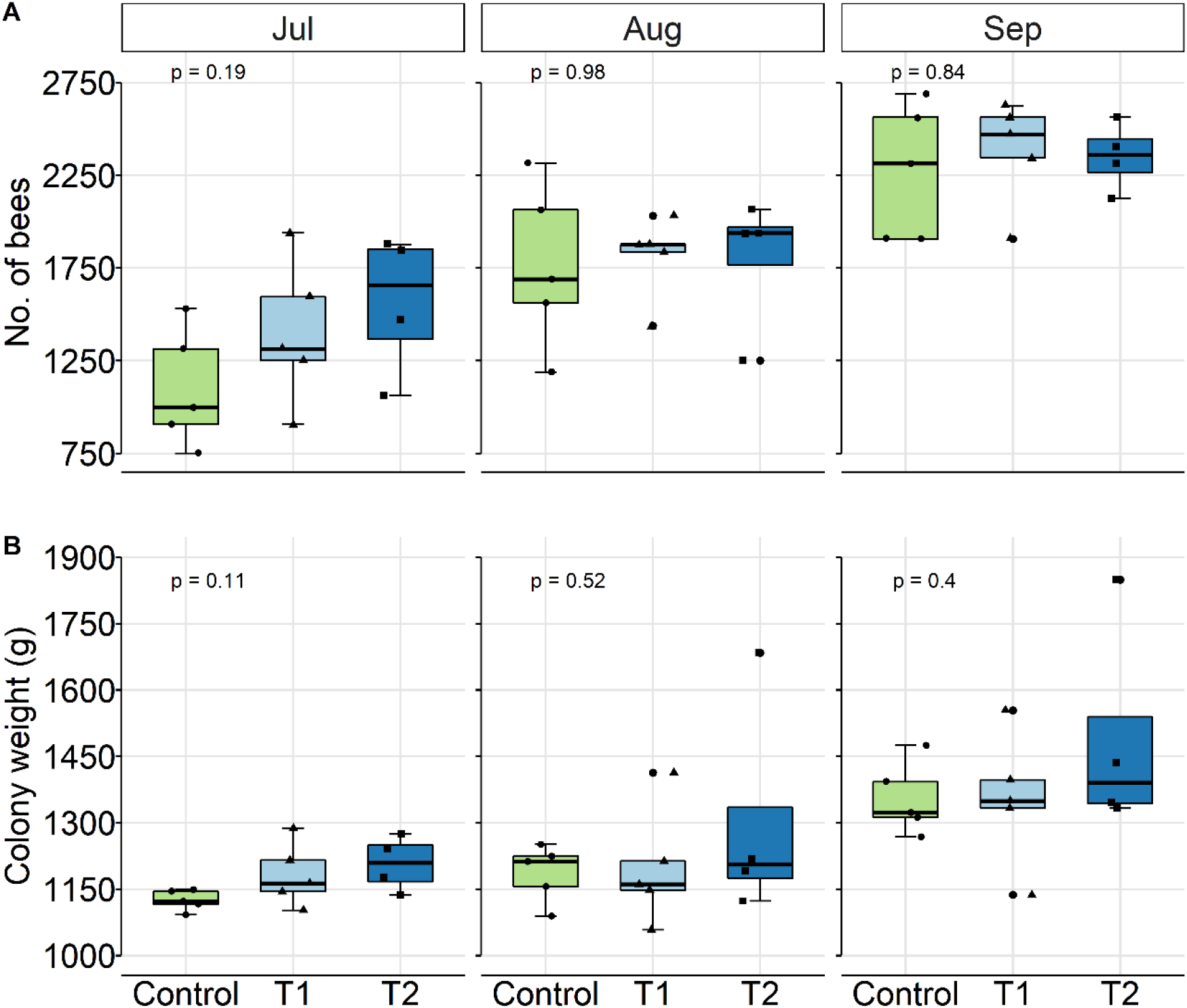
**A** Boxplot displaying the number of worker bees and their increase during the course of the study in the different treatments (control C, glyphosate treatment T1: 4.8 mg a.i./kg, T2: 137.6 mg a.i./kg). **3B** Boxplot displaying the absolute colony weight and its increase during the course of the study. There were no significant differences between groups (*P* > 0.05, ANOVA)]

**Fig. 3.**
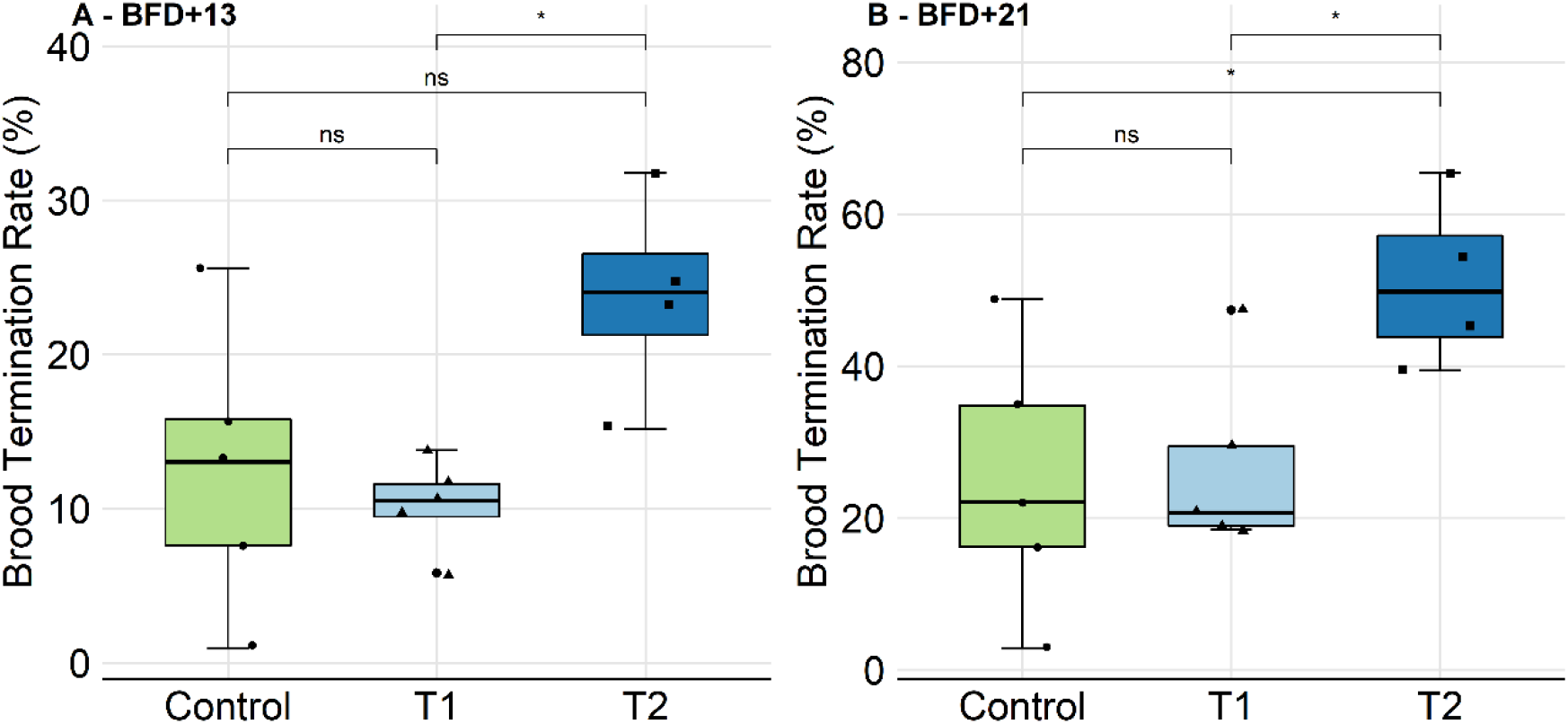
**A** Brood Termination Rate (BTR) from BFD+13 in%. Treatments T1 and T2 did not show differences when compared to C (*P* > 0.05, t-test, pairwise). **4B** BTR from BFD+21. While T1 does not show differences when compared to C (*P* > 0.05, t-test, pairwise), T2 revealed a significantly higher BTR (*P* < 0.05, t-test, pairwise).]

#### 3.1.2 Brood development and photographic assessment

With reference to the developing time of a worker honey bee from egg to adult (21 to ±1 days (Jay 1963, Wang et al. 2015)), it was assumed that all eggs should completely be developed at the time of the last assessment date (BFD+21). The cumulative number of selected eggs from all replicates was C (n=522), T1 (n=525) and T2 (n=426) with no significant difference of the mean distribution amongst groups C (n=104/replicate), T1 (n=105/replicate) and T2 (n=106/replicate) (Fig. S3, *P* > 0.05, ANOVA).

In groups C, T1 and T2, successful development was observed in the majority of the marked brood cells. On BFD+13, the median termination rate was 13.04/10.53%, respectively. In the test item treatment T2, brood termination was not significantly increased at this point, when compared to the control (24.05%, Fig. 4A).

**Fig. 4.**
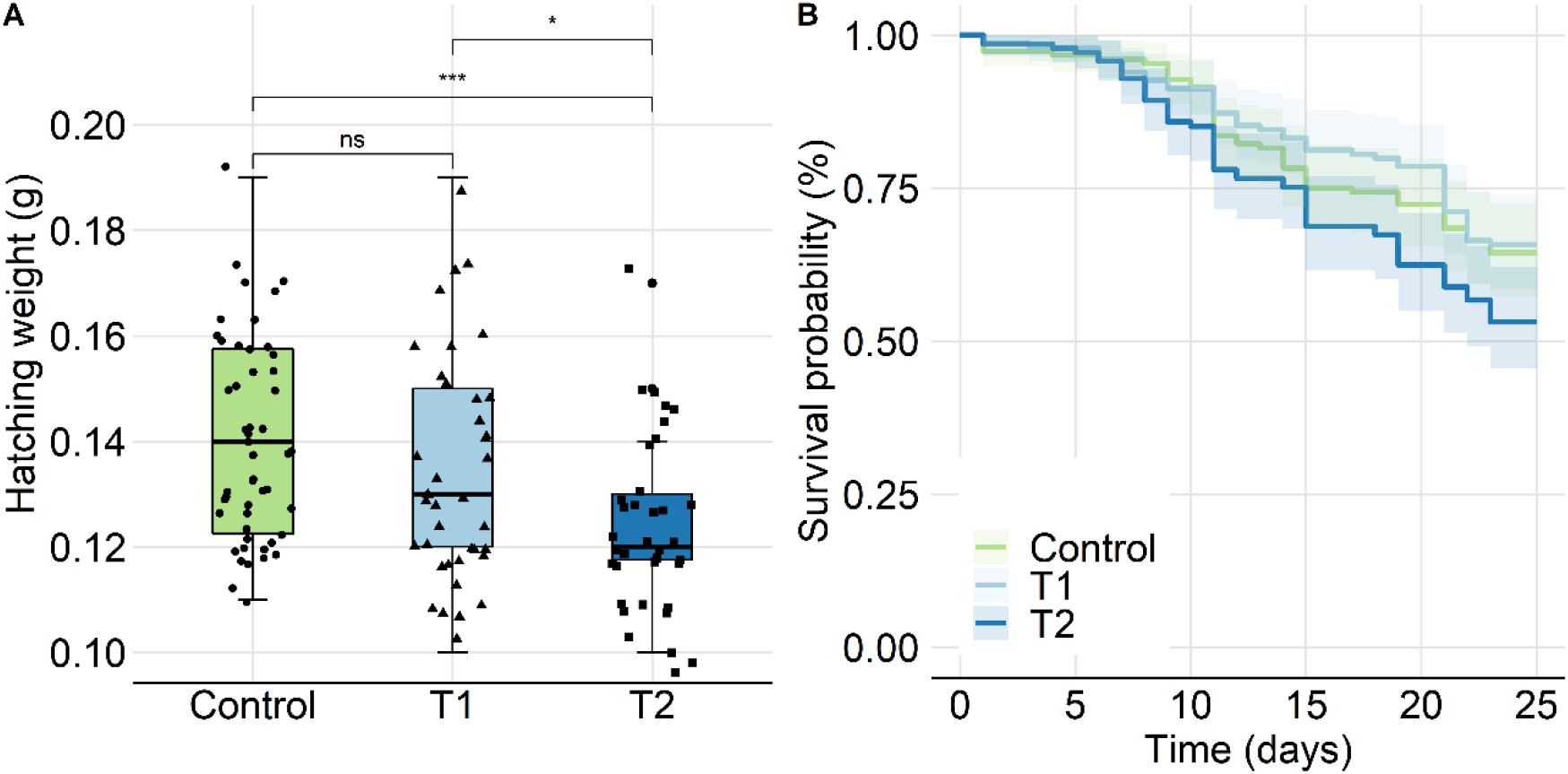
**A** Hatching weight assessed on the day of emergence after 24 h in the incubator (see Fig. 2). Weight in T2 was significantly lower when compared to C (*P* < 0.01, t-test) and T1 (*P* < 0.05, t-test). **5B** Groups were compared with a Kaplan-Meier-Survival analysis (control C, glyphosate treatment T1: 4.8 mg a.i./kg, T2: 137.6 mg a.i./kg). Survival probability is displayed with 95% confidence intervals. Significant differences within those groups were revealed close to the statistical threshold (global *P* = 0.047, log-rank test). A pairwise comparison, however, did not confirm these differences between the respective groups (*P* > 0.05, log-rank test).]

On BFD+21, the median termination rate was 22.11/20.69%, respectively. In the test item treatment T2, brood termination was significantly increased when compared to the control (*P* < 0.05, t-test, pairwise). The median termination rate was 49.84% (Fig. 4B). Full detail of the brood assessments is provided with the supplementary material (Fig. S4 and Method S1).

#### 3.1.3 Survival of individual bees and hatching weight

The hatching weight of adult workers in T2 was significantly lower when compared to C (*P* < 0.01, t-test) and T1 (*P* < 0.05, t-test) with median values of C: 0.14, T1: 0.13 and T2: 0.12 g (Fig. 5A). Further, the Kaplan-Meier-Survival analysis of all three groups indicated a significant difference (Fig. 5B, global *P* = 0.047, log-rank test). When compared pairwise, however, no significant differences were confirmed between the respective groups (*P* > 0.05, log-rank test). In addition, a Cox proportional hazards model was applied to determine the hazard ratio (HR) displayed as forest plot (Fig. S5). With an HR of 0.93 for T1 and 1.43 for T2, the treated bees were not at risk of dying sooner when compared to the control (T1: *P* = 0.73, T2: *P* = 0.051, log-rank test). To justify pooling bees from the same groups but different mini-hives for survival analysis, these hives were evaluated separately treated as replicates. The test showed no significant differences (Fig. S6, *P* > 0.05, log-rank test).

**Fig. 5.**
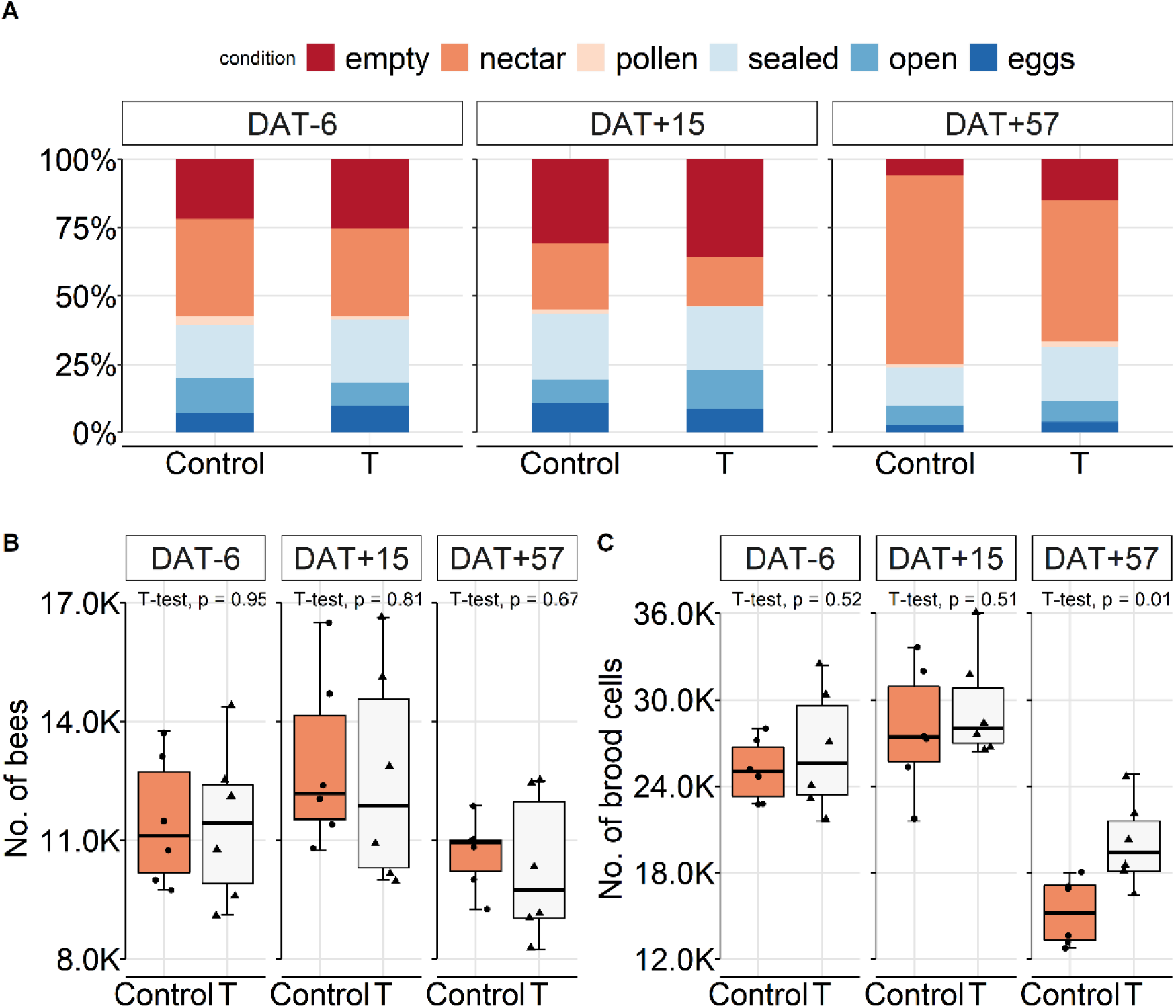
A Proportion of empty cells, stores (nectar, pollen) and worker brood cells (eggs, open, sealed) presented as total colony condition of both groups (control C, glyphosate treatment T) corresponding to their assessment date (DAT = day after treatment). 6B/C Boxplot displaying the number of worker bees/brood cells during the course of the study. Dates are corresponding to the following months; DAT-6 (=Jul), DAT+15 (=Aug), and DAT+57 (=Sep).]

### 3.2 Field exposure and overwintering

#### 3.2.1 Colony conditions

Results from 2.2 (experiment 2) and 2.3 (experiment 3) are presented in a combined approach (Fig. 6 and Fig. 7, respectively). Overall, colonies were well provided and did not suffer food shortage. GBH did not significantly affect colony development in both experiments, regardless of the type of exposure and time of the year (*P* > 0.05, t-test, pairwise). In experiment 2, bees and brood showed a pattern of increases and decreases in an alternate sequence (Fig. 6B/C). However, brood cells were decreasing faster in C on DAT+57 due to a greater winter food intake (Fig. 6C, *P* = 0.014, t-test).

**Fig. 6.**
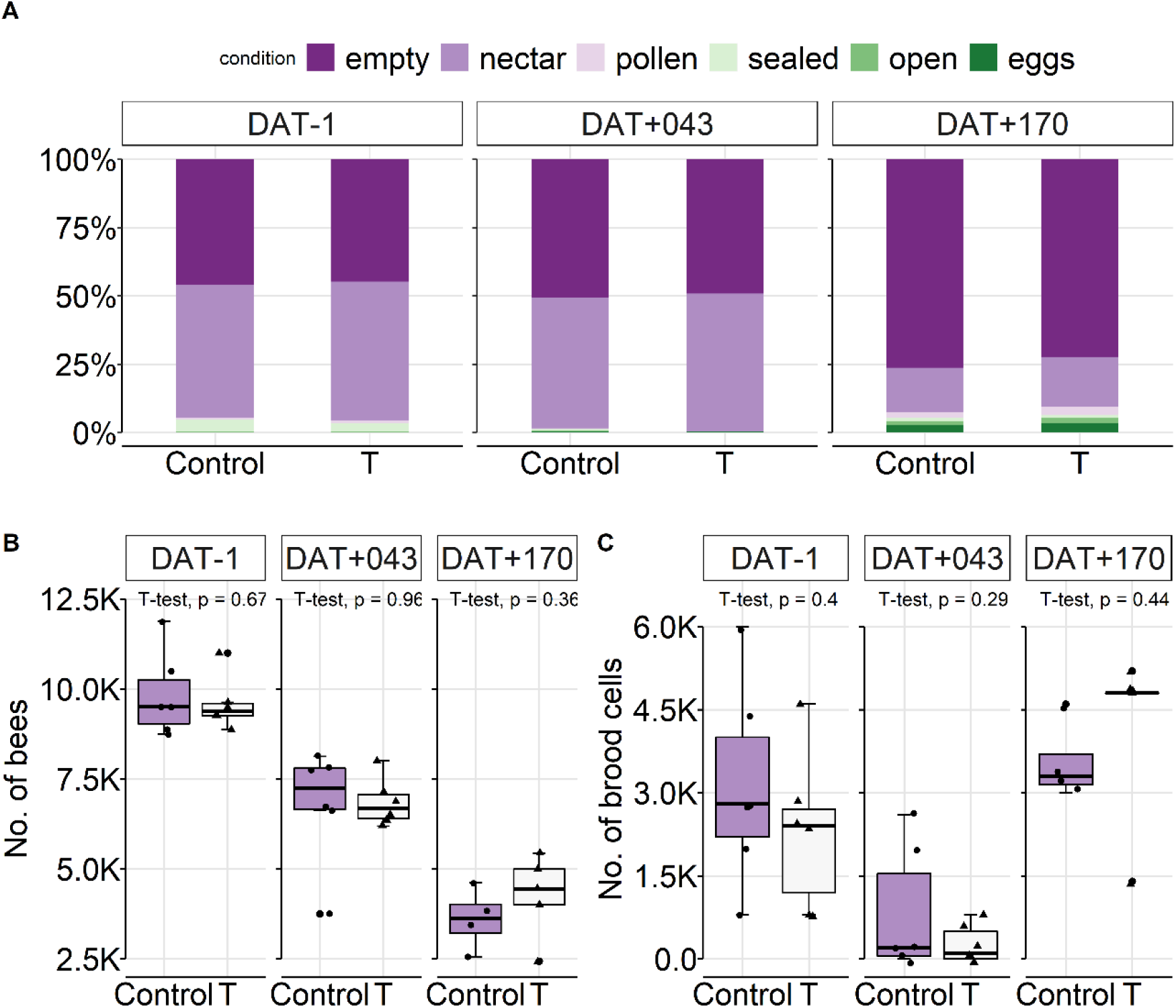
**A** Proportion of empty cells, stores (nectar, pollen) and worker brood cells (eggs, open, sealed) presented as total colony condition of both groups (control C, glyphosate treatment T) corresponding to their assessment date (DAT = day after treatment). **7B/C** Boxplot displaying the number of worker bees/brood cells during the course of the study. Dates are corresponding to the following months; DAT-1 (=Oct), DAT+43 (=Nov), and DAT+170 (=Mar).]

In experiment 3, shortly before hibernation (DAT+43), brood activity was reduced accordingly (Fig. 7C) followed by a continuous decrease of bees (Fig. 7B). At the end of overwintering, a decrease of nectar (food) was observed (Fig. 7A) as a result of the returning brood activity (Fig. 7C). On DAT+170, we found two colonies in C and one in T having not survived winter.

### 3.3 Residues

The results of the residue analyses for glyphosate and its metabolite aminomethylphosphonic acid (AMPA) from all four experiments are shown in Tab. 2. A total of 176 samples were measured including controls (experiment 1: 6, experiment 2: 92, experiment 3: 66, experiment 4: 12) (see Odemer, 2020b). It could be confirmed that all experimental colonies from the respective treatments were exposed to glyphosate, accordingly.

In experiment 1, 2 and 3, irrespective of the time interval after the application (short term: three days or long term: 170 days), glyphosate residues remained constant in all sugar matrices. In experiment 2 a 15.9-fold average increase, and a 24-fold peak increase (see raw data) was measured in stored pollen (beebread) when compared to stored nectar. In plants, a range similar to experiment 4 (tunnels) was measured shortly after the application on DAT0. Degradation of glyphosate in the tunnels, however, was faster than in the field. In experiment 4, a matrix dependent exposure gradient could be identified, which can be presented from high to low glyphosate residues as follows: corbicular pollen > plants > honey sac > stored food/nectar. Honey sac residues were measured with a 3.3-fold reduction when compared to their floral source (plants). In turn, a 27.4-fold increase was measured in pollen when compared to honey sac contents, similar as reported in experiment 2 for the stored products.

Notably, in experiment 1 trace residues of glyphosate were found in the pooled control food on DAT+21 (0.18 mg a.i./kg, AMPA was not detectable). Further, in experiment 3, trace residues of glyphosate were found in one control and one treatment colony before the application on DAT-1 (C2: 0.05 and T2: 0.04 mg a.i./kg, AMPA was not detectable).

## 4 DISCUSSION

Honey bee workers in our study that were chronically exposed to GBH in a sensitive stage of development (larval and pupal phase) did not die sooner in reference to the untreated bees. Colonies also did not break down during overwintering nor were affected when directly foraging on GBH sprayed crops, at least not with the here used formulated products. Their development followed a typical pattern considered normal for a temperate climate (Imdorf et al. 2008). To our surprise, the termination rate (BTR) was significantly increased on BFD+21 after the brood was chronically exposed to the high concentration (T2) in the feeding phase. A trend became evident at BFD+13, already. This is particularly interesting as the number of bees and the colony weight did not decline but increased during the course of the study, despite the almost doubled BTR in T2. Our finding is supported by Vazquez et al. (2018) who reported similar effects when larvae were exposed to chronic GBH feeding *in vitro*, as they state that molting was delayed and fresh weight was reduced. The hatching weight of young bees in our study was significantly reduced in the high concentration (T2), too. With a median of 120 mg, however, it remains within the variation of reported fresh weight in the literature. Kunert and Crailsheim (1988) found the mean fresh weight of summer bees to be 116.7 mg (± 1.5 SE), where Żółtowska et al. (2011) state a range from 86.08 mg to 121.18 mg. In contrast, Thompson et al. (2014) found larvae/pupae not affected, nor was there a negative impact on pupal weight using the feeding method described by Oomen et al. 1992. Therefore, brood assessments were only conducted until day 16 of development, which does not fully cover a complete worker brood cycle. Moreover, it needs to be emphasized that for the brood study technical grade isopropylamine (IPA) salt was used instead of a formulated glyphosate product. This can be of relevance concerning the differences in the outcome of our and Vazquez’s study (Vazquez et al. 2018) and suggests that the IPA salt alone may not be the striking element accountable for brood effects. Previously, evidence has emerged that adjuvants found in formulations of plant protection products may not be inert as alleged, but can be toxic to bees (Mullin et al. 2016). As these adjuvants are currently regulated differently from active ingredients, it cannot be ruled out that the here observed discrepancy derives from the presence of such substances (Mesnage et al. 2012, Mesnage and Antoniou 2018). Thompson et al. (2014) further assessed colony conditions with the “Liebefeld method” (Imdorf et al. 1987) and obtained results that are in line with those of our field studies. No negative effects on colony growth and overwintering success could be revealed. These findings overall suggest that glyphosate does not have a lasting effect on honey bee population dynamics, but is able to delay pre-adult development of worker bees as a formulated product.

Following the method of Schur et al. (2003) only one single brood cycle was covered in the mini-hive experiment. Thus, information on the extent pupation was delayed by GBH are currently lacking. We suggest that, in a next step, evaluation of at least a second brood cycle is required to provide such details. This knowledge would allow us to further assess possible colony-level consequences of GBH. Most notably, a delay in brood development may result in an unintentional reproductive advantage for the mite *V. destructor*. Its reproductive cycle is precisely aligned with the host brood development (Kirrane et al. 2012, Frey et al. 2013). The viable offspring generated within one mite cycle in worker brood with regular development (21 days) is measured at 1.1 to 1.4 daughter mites per foundress (Fuchs und Langenbach 1989, Donze et al. 1996). The consequence of a delayed worker brood development could possibly lead to a higher rate of viable daughter mites to be released from infested cells, boosting *Varroa* numbers within colonies.

Retrospectively, there also seems to be a contradictory terminology in the method we followed, and which is used in regulatory risk assessment to evaluate the effects of plant protection products on honey bees (Schur et al. 2003, OECD 2007, Oomen et al. 1992). The term “Brood Termination” implies that development was ended or stopped in certain cells because the expected brood stage was not appropriately achieved (Tab. 1). A delay, however, is not reflected correctly by this term. Hence, we suggest to improve the method and refine its terminology as there have emerged new insights to limit the susceptibility to errors, recently (Wang et al. 2020). Especially under semi-field conditions, high BTR’s in control colonies are an issue (Lückmann et al. 2015) that are causing high variability within replicates. Suggestions by the ICP-BR Bee Protection Group resulted in improvements of the method, however, sublethal effects were not in focus (Pistorius et al. 2012). With current technical advances it is possible to enhance the selection process of study colonies, digitize colony and brood assessments and monitor flight activity to display sublethal effects more accurately (Colin et al. 2018, Bermig et al. 2020, Wang et al. 2020). This refinement could be of high relevance to further improve regulatory relevant methodologies to identify sublethal effects at the colony level and help to render present terminology more precisely.

**Tab. 1.** Glyphosate (GLY) and aminomethylphosphonic acid (AMPA) residues corresponding to the assessment date (DAT = day after treatment) and matrix from all four experiments. For the detection limits of the method see materials and methods section. All values are rounded and presented as means (± SD) in mg a.i./kg where appropriate (for details of the method and repeat determination, see supplementary material Method S2 and raw data Odemer, 2020b. *: pooled sample, +: analyzed twice, n.d.: not detectable, −: not measured)]

|  | Time | Samples (n) | Stock solution |  | Stored food/nectar |  | Honey sac |  | Beebread |  | Corbicular pollen |  | Plants |  |
| --- | --- | --- | --- | --- | --- | --- | --- | --- | --- | --- | --- | --- | --- | --- |
|  |  |  | GLY | AMPA | GLY | AMPA | GLY | AMPA | GLY | AMPA | GLY | AMPA | GLY | AMPA |
| Experiment 1 | DAT0 (T1) | 1 | 4.764 ± 0.424<sup>+</sup> | 0.020 ± 0.003<sup>+</sup> | - | - | - | - | - | - | - | - | - | - |
| | DAT0 (T2) | 1 | 137.566 ± 4.318<sup>+</sup> | 0.428 ± 0.001<sup>+</sup> | - | - | - | - | - | - | - | - | - | - |
|  | DAT+21 (T1) | 5* | - | - | 5.103 | 0.013 | - | - | - | - | - | - | - | - |
|  | DAT+21 (T2) | 4* | - | - | 99.861 | 0.319 | - | - | - | - | - | - | - | - |
| Experiment 2 | DAT0 (+0.5h) | 6* | - | - | n.d. | n.d. | - | - | - | - | - | - | 96.429 ± 13.540<sup>+</sup> | 0.360 ± 0.005<sup>+</sup> |
| | DAT+1 | 6 | - | - | 0.327 ± 0.246 | 0.087 ± 0.194 | - | - | - | - | - | - | 44.715 ± 1.429<sup>+</sup> | 0.255 ± 0.007<sup>+</sup> |
| | DAT+3 | 6 | - | - | 0.490 ± 0.455 | n.d. | - | - | - | - | - | - | 33.634 ± 0.016<sup>+</sup> | 0.218 ± 0.017<sup>+</sup> |
| | DAT+6 | 6 | - | - | 0.347 ± 0.362 | n.d. | - | - | 5.512 ± 6.922 | 0.007 ± 0.017 | - | - | - | - |
| | DAT+7 | 6 | - | - | - | - | - | - | - | - | - | - | 31.051 ± 1.930<sup>+</sup> | 0.224 ± 0.007<sup>+</sup> |
| Experiment 3 | DAT-1 | 12 | - | - | 0.004 ± 0.012 | n.d. | - | - | - | - | - | - | - | - |
| | DAT+6 | 6 | - | - | 4.462 ± 1.651 | 0.006 ± 0.005 | - | - | - | - | - | - | - | - |
| | DAT+13 | 6 | - | - | 5.159 ± 1.011 | 0.012 ± 0.004 | - | - | - | - | - | - | - | - |
| | DAT+43 | 6 | - | - | 7.148 ± 3.234 | 0.033 ± 0.015 | - | - | - | - | - | - | - | - |
| | DAT+170 | 6 | - | - | 4.985 ± 0.613 | 0.019 ± 0.004 | - | - | - | - | - | - | - | - |
| Experiment 4 | DAT0 (+1h) | 3* | - | - | - | - | 24.918 | 0.092 | - | - | 614.841 | 2.122 | 82.407 | 0.217 |
|  | DAT+1 | 3* | - | - | - | - | - | - | - | - | - | - | 103.892 | 0.362 |
|  | DAT+2 | 3* | - | - | - | - | - | - | - | - | - | - | 62.845 | 0.229 |
|  | DAT+3 | 3* | - | - | - | - | - | - | - | - | - | - | 92.886 | 0.395 |
| | DAT+5 | 4 | - | - | 0.182 ± 0.225 | n.d. | - | - | - | - | - | - | - | - |

One particular sublethal effect that recently gave rise to concerns was induced by GBH (Motta et al. 2018). Here, glyphosate altered abundances of bacterial communities when exposed to field-realistic concentrations and further increased susceptibility of bees to the opportunistic pathogen *Serratia marcescens*, which can be lethal under certain conditions (Raymann et al. 2018). In this context, the findings of Zheng et al. (2017) should be highlighted. The authors provide evidence, that gut bacteria influence metabolic functions of their host with a potential contribution to host nutrition - affecting weight gain in particular. As suggested by our results and the results of Vazquez et al. (2018), high GBH exposure obviously had an effect on larval and adult fresh weight. Presumably caused by a dysfunction of metabolic processes, this should be further investigated. Specifically designed experiments taking into account realistic exposure scenarios of GBH could reveal possible consequences at the colony level.

In the current debate about the potential risk for pollinators deriving from GBH, critical voices have been raised to immediately ban glyphosate from arable fields and home use (European Commission, 2020). To date, it is, however, largely unknown to which scale glyphosate residues are present in the environment of pollinators, especially in cultivated plants attractive to bees. Due to the complex analysis of glyphosate, it cannot be integrated in a multi-method (Székács and Darvas 2012). Thus, GBH are not subject to many routine assessments of standard bee matrices such as honey and pollen. In this study, we wanted to complement existing residue data. We revealed that glyphosate residues in pollen are up to 27-fold higher when compared to residues in nectar (collected fresh and from stored in-hive products), deriving from the same application. Our finding is consistent with that of Thompson et al. (2014) who presented the same order of magnitude in their green-house experiments. Here, glyphosate values were approximately 18-fold higher in pollen as compared to nectar. This deviation cannot be attributed to chemical properties of glyphosate, the ammonium salt is hydrophilic and has an amphoteric (dipolar ionic) character (Székács and Darvas 2012). The epicuticular wax on the surface of plant leaves represents a barrier to the polar properties of glyphosate. Hence, so-called surfactants are added to GBH formulated products to overcome this barrier and increase efficacy (Curran and Lingenfelder 2009). We assume that due to this fact, pollen shows a many-fold increase in residue levels towards nectar, as the surfactant enriches the active ingredient in the lipophilic matrix of pollen grains. Further, we are in line with the results provided by Karise et al. 2017 and Rubio et al. 2014 but contradicting Berg et al. 2018, who found many-fold higher glyphosate concentrations in honey than we and the others did. It seems reasonable, that in this particular study honey bees must have been exposed to unusually high levels of GBH, which could be a result of misapplication or misuse of the product. However, it seems likely that free-foraging bees are at continuous risk of contamination. In experiment 1 glyphosate residues of 0.18 mg/kg have been measured in the pooled control sample from DAT+21. Due to the setup of the mini-hives, drifting of bees was possible and, even if not obviously visible, robbing could not be ruled out. Notably, in experiment 3 one control and one treatment colony had residues of 0.05 and 0.04 mg glyphosate/kg, respectively, before the application. This indicates that glyphosate was most likely introduced from the environment. However, after seven days, residues were no longer present in the control. From these low numbers we do not assume side-effects compromising the outcome of our study, but we suggest to take measures preventing unintentional contamination in future experiments. A less densely placement of the hives would be a good start to reduce drifting.

Our results further provide a baseline of residues that can be found in a realistic exposure scenario. We point to the fact that glyphosate was persistent up to 170 days in sugar-based matrices such as honey and syrup. Microbial degradation of glyphosate is prominent for a wide range of gram-positive and negative bacteria, and for different classes of fungi (reviewed in Sviridov et al. 2015). Interestingly, honey is long-known for its antimicrobial qualities (Israili 2014). Its high sugar contents result in a low water activity (*a*_*w*_), which inhibits or stresses bacterial growth (Chirife et al. 1983). Due to this characteristic, we suggest that sugar-based matrices are a glyphosate-friendly environment that prevents degradation of the active ingredient. Moreover, glyphosate degraded more quickly in phacelia plants grown under field conditions compared to the tunnel plants. This can most probably be attributed to oxidative advanced processes such as photolysis due to ultraviolet (UV) light exposure (Assalin et al. 2009). The tunnel gauze obviously protected the active ingredient from a faster degradation, possibly increasing the action of the herbicide. This is particularly interesting when considering more harmful substances such as insecticides and should be investigated in more detail. A tunnel exposure, therefore, very likely represents a worst-case scenario where bees are not only challenged to the confinement but also to the limited and unbalanced forage (OECD 2007, EPPO 2010). This outcome can be helpful to categorize upcoming findings and provide support to future experiments.

When investigating plant protection products, recent attention has mostly been drawn to neonicotinoid insecticides. Fungicides and herbicides are currently understudied substance classes in a bee health context (Havard et al. 2019, Cullen et al. 2019). With regard to our findings, however, we find they deserve closer attention. As a lesson learned from past “neonicotinoid research” a clear focus on field level data should be emphasized for upcoming studies. We could see that many negative effects and various synergisms found in individuals at the laboratory level did not necessarily translate to negative effects in the field at the colony level (Retschnig et al. 2015, Sponsler & Johnson 2017). Paris et al. 2020 reported, for instance, that the fungicide boscalid commonly applied to oilseed rape, acts in synergy with the gut parasite *Nosema ceranae* under laboratory conditions. The authors found that microbiota were disturbed by the fungicide and suggested that bacterial metabolism was disrupted analog to glyphosate effects reported in Motta et al. (2018). This led to a significantly higher mortality in combination with *N. ceranae* when compared to boscalid exposure or parasite infection alone. It is, however, unclear how and whether this effect can be extrapolated to the colony level. Hence, field data generated under beekeeping conditions would be far more conclusive taking bee research to the necessary next step. In another example, this was convincingly demonstrated by Wernecke et al. (2019). The authors found that ergosterol biosynthesis-inhibiting (EBI) fungicides combined with pyrethroids or neonicotinoids can lead to a synergistic toxic increase affecting the survival rate of honey bees under laboratory, semi-field and field conditions – covering many stages of effects from laboratory to field. We therefore suggest that future research should not only focus on the above-mentioned understudied substance classes but should also implement holistic testing regimes. Investigating possible synergistic effects with both, parasites and tank-mixtures applied in the field could capture a realistic impression of the honey bee health status. Provided that test systems consider field realistic exposure scenarios under beekeeping conditions with higher emphasis.

## 5 CONCLUSIONS

Glyphosate-based herbicides are considered not harmful to honey bees but can be found all over conventional agro-ecosystems worldwide. Many apiaries are in close proximity to agricultural landscapes and bees are at risk of foraging on potentially contaminated food sources. Even though we confirmed that glyphosate does not have an impact on the lifespan of individuals, colony conditions and overwintering, it became obvious that the herbicide delayed worker brood development when applied as a chronic high dose. Our results corroborate findings from the laboratory, where a delay in larval development was reported. In addition, GBH exposure resulted in reduced body weight of exposed individuals in both, laboratory and our field study. However, when considering that stored nectar and beebread are typically utilized for brood rearing, our results need to be interpreted with caution. Bearing in mind the low residues we found in these two matrices, negative effects at the colony level are not likely to be expected when GBH are applied under good agricultural practice. Regardless, we highlight the importance of suitable and relevant approaches to detect the slightest sublethal effect possible, which is not necessarily provided with current standard regulatory methods. The refinement of existing methodology could reveal lasting consequences not only for honey bee but also pollinator health, which we are perhaps not able to observe at present. We therefore see a great need to improve such implementations in future research. Technological advances such as automated bee counters or hive scales could help detecting sublethal effects more easily. Our present data will in addition greatly contribute to the existing evaluation of field-realistic residues that are to be expected from GBH exposed bee attracting crops. Necessarily, further studies are needed to complete the big picture of a realistic GBH exposure scenario to which not only honey bees but other pollinators are challenged under field conditions. The evidence provided here could serve as basis for future experiments to narrow down the exposure range bees are confronted within the environment. Ultimately a pan-European pesticide monitoring could be established, where such results are merged into a database that serves as information for the public, beekeepers, and decision-makers.

## ACKNOWLEDGEMENTS

We appreciate the support of our technical staff for the excellent laboratory and field-work, in particular we want to thank Benjamin Grasz and Charlotte Steinigeweg for their help. Further, we are grateful to Silvio Erler for his review and provision of valuable comments on an earlier draft of the manuscript.

This research did not receive any specific grant from funding agencies in the public, commercial, or not-for-profit sectors.

The data that support the findings of this study are available in the “Open Science Framework” (Odemer, 2020b).

## Ethical approval

This article does not contain any studies with human participants or animals performed by any of the authors.

## Author contribution

Conceptualization, R.O., F.O., I.W., A.W. and M.F.; Methodology, R.O., I.W., and M.F.; Software, R.O.; Validation, R.O., G.B. and A.A.; Formal Analysis, R.O., G.B.; Investigation, R.O., F.O., A.A., A.W., G.B. and M.F.; Resources, R.O., F.O., J.P.; Data Curation, R.O.; Writing – Original Draft Preparation, R.O.; Writing – Review & Editing, A.A., G.B., M.F., J.P., I.W., A.W., F.O.; Visualization, R.O.; Supervision, J.P., R.O., G.B.; Project Administration, R.O., I.W., A.W. and M.F; Funding Acquisition, R.O., F.O. and J.P.

