## Supplementary Material for "Chronic high glyphosate exposure delays individual worker bee (*Apis mellifera* L.) development under field conditions"

Tab. S1: Color code of labelled cell contents (BFD)

Fig. S1: Pictures of combs with color code

Fig. S2: Experimental setup scheme (experiment 1)

Fig. S3: Selected mean No. of eggs

Fig. S4: BTR, Brood Index and Compensation Index from BFD+13 and BFD+21

Fig. S5: Hazard ratio (HR) displayed as forest plot

Fig. S6: Survival probability of untreated mini-hive replicates to justify pooling

Method S1: Brood development and photographic assessment adopted from OECD 2007 and based on Schur et al. 2003

Method S2: Analytical method and validation for glyphosate and AMPA

### **Supplementary Table S1:** Color code of labelled cell contents (BFD)

**Tab. S1** Brood area Fixing Day (BFD) assessments during the course of the chronic glyphosate exposure, modified after Schur et al. (2003). The numbers for the brood index were assigned to the cell content, respectively. If the expected brood stage was met at the specified BFD, the cell was labeled “1”, if not as terminated (Brood termination = 0). To differentiate brood stages in the photographic assessment, a color code was used (see Fig. S1).

| Timing | Expected brood stage | Brood index | Brood termination | Color code |
|---|---|---|---|---|
| BFD0 | egg | 1 | - | blue |
| BFD+6* | young (L1-L2) to old larva (L3-L5) | 2, 3 | if ≠ 2,3 | green, red |
| BFD+13 | capped cell (pupa) | 4 | if ≠ 4 | purple |
| BFD+17* | capped cell (pupa) | 4 | if ≠ 4 | purple |
| BFD+21 | empty cell (e), egg, young larva, nectar (n), pollen (p) | 5 | if ≠ 1, 2, e, n, p | yellow |

**Supplementary Figure S1:** Pictures of combs with color code

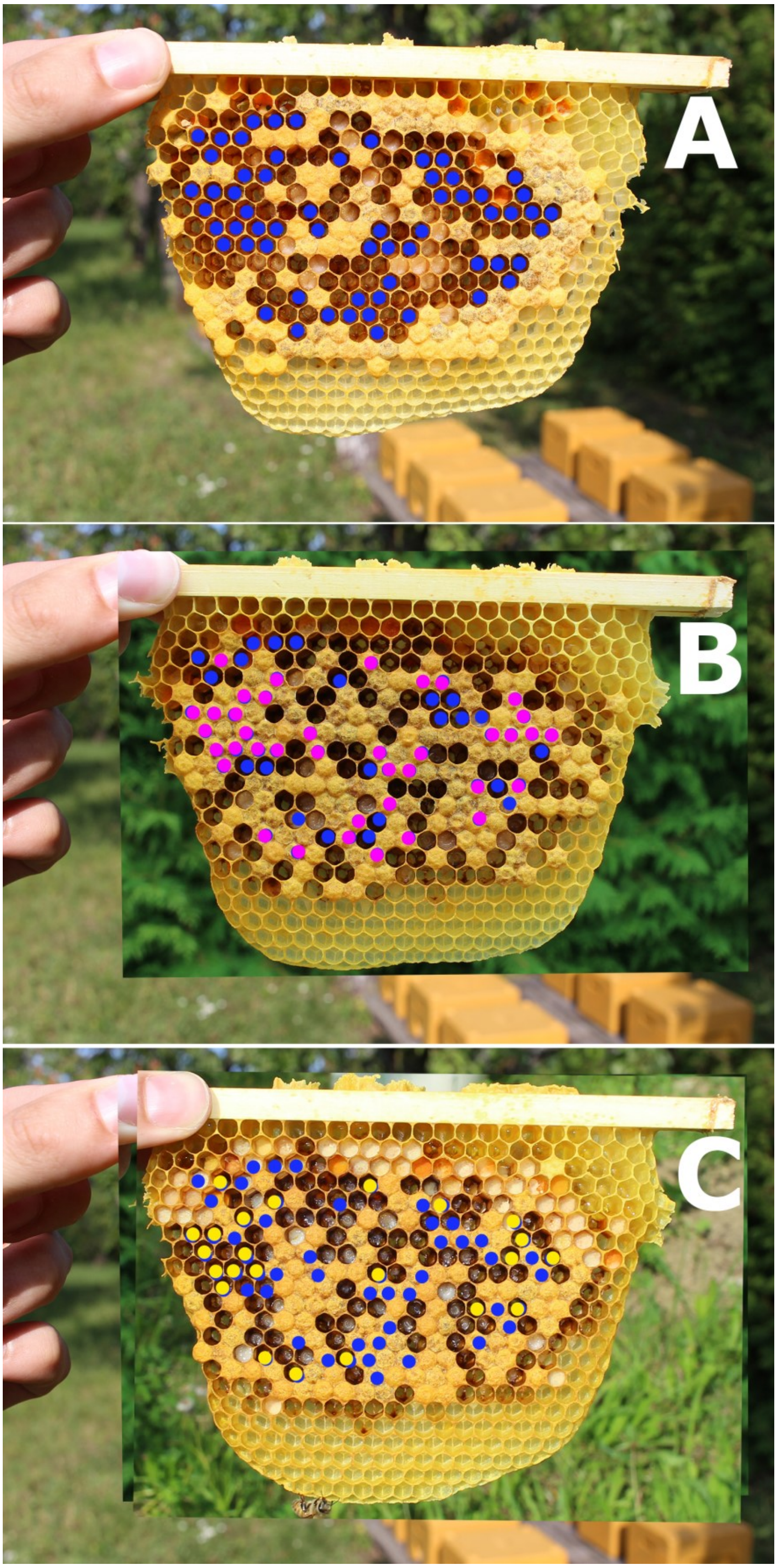

**Fig. S1** Pictures from comb no. 4, side A, colony V12, treatment T2. A color code was applied to identify different developmental stages of the brood according to Tab. 1. **A** BFD0, eggs, blue, **B** BFD+13, capped cells (pupae), purple, **C** BFD+21, empty cells OR eggs OR pollen OR nectar OR young larvae, yellow. With this color code, cells could be identified where development was successfully achieved according to Tab. 1. In the case where this was not the case, the development was regarded as terminated. For BFD+13 and BFD+21 the cumulative brood termination rate, brood index and compensation index were calculated for each colony (Fig. S3, Method S1).

### **Supplementary Figure S2:** Experimental setup scheme (experiment 1)

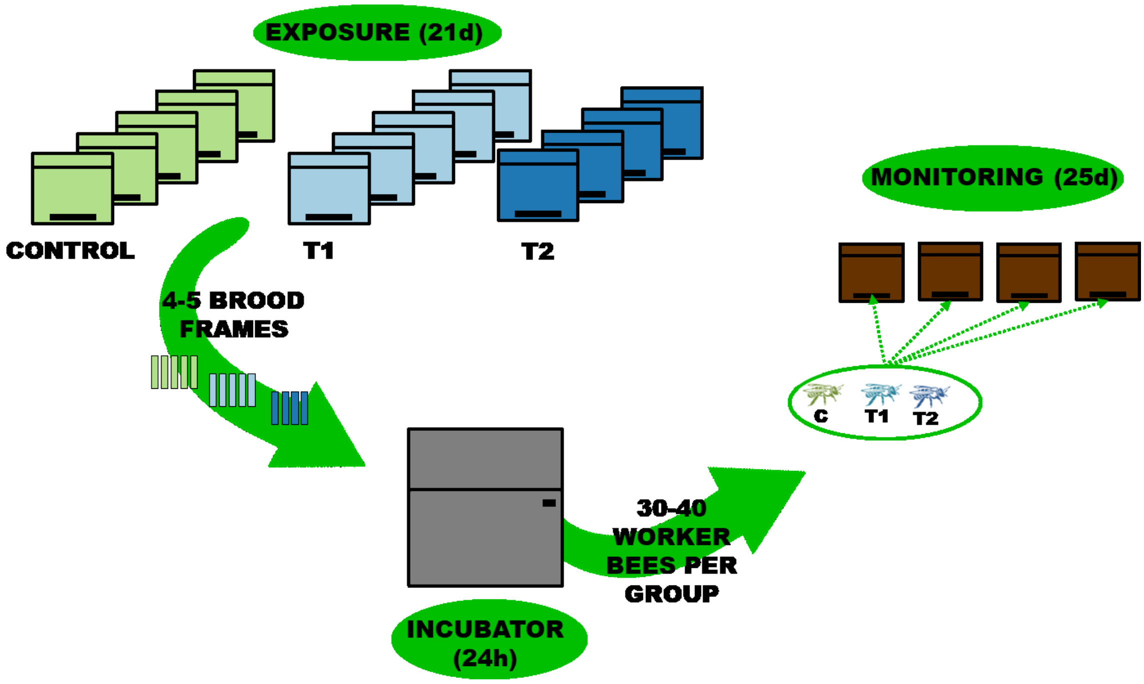


**Fig. S2** Experimental setup illustrating the exposure and monitoring period. Control and T1 were comprised of five mini-hives, T2 four. Colonies were exposed to glyphosate for 26 days. Subsequently, one ready-to-hatch brood frame per mini-hive was removed and placed together group-wise in an incubator for 24 h. After hatching, a total of C (n=152), T1 (n=149) and T2 (n=141) bees were marked and subdivided. Approximately 30-40 worker bees per treatment were introduced into four untreated mini-hives, each, and monitored for 25 d.

**<u>Supplementary Figure S3:</u>** Selected mean No. of eggs

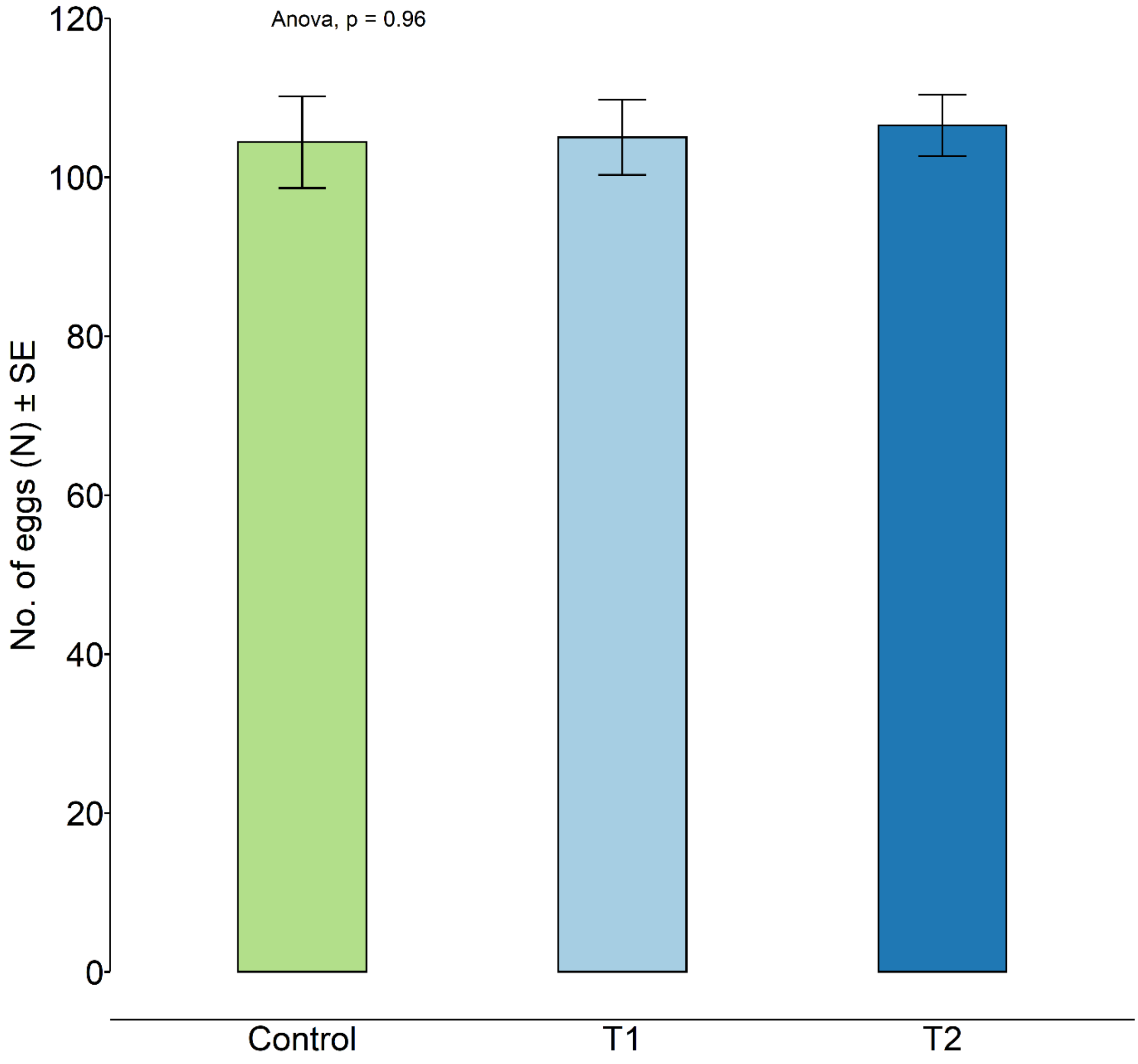


**Fig. S3** Barplot displaying mean number of selected eggs for the digital brood assessment on BFD0. All replicates had a similar number of selected eggs (approximately 100).

**Supplementary Figure S4:** BTR, Brood Index and Compensation Index from BFD+13 and BFD+21

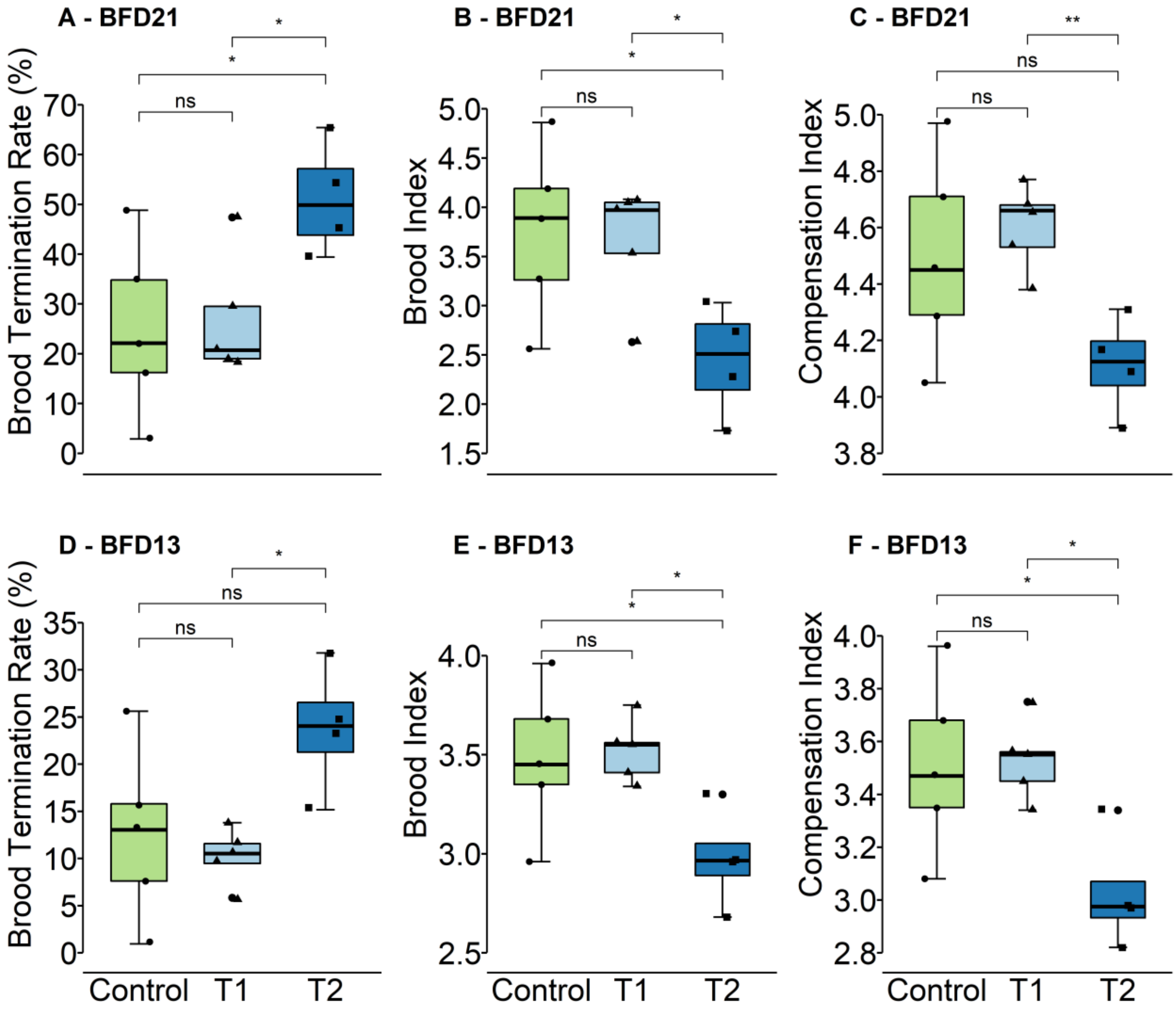


**Fig. S4** Here, full details from the brood assessment are presented modified after Schur et al. (2003) to complement Fig. 4 (see also Method S1). In groups C and T1, successful development was observed in the majority of the marked brood cells. Assuming that at the first assessment only eggs will be marked, the index is 1.0. An increase of the brood index (see paragraph 40) during the following assessment can be observed, if a normal development of the brood is presumed. This increase is caused by the development from eggs to larval stages, to the pupae and finally to the adult, emerged bee and furthermore due to the rising numbers which are assigned to the brood stages (OECD, 2007).

**Supplementary Figure S5:** Hazard ratio (HR) displayed as forest plot

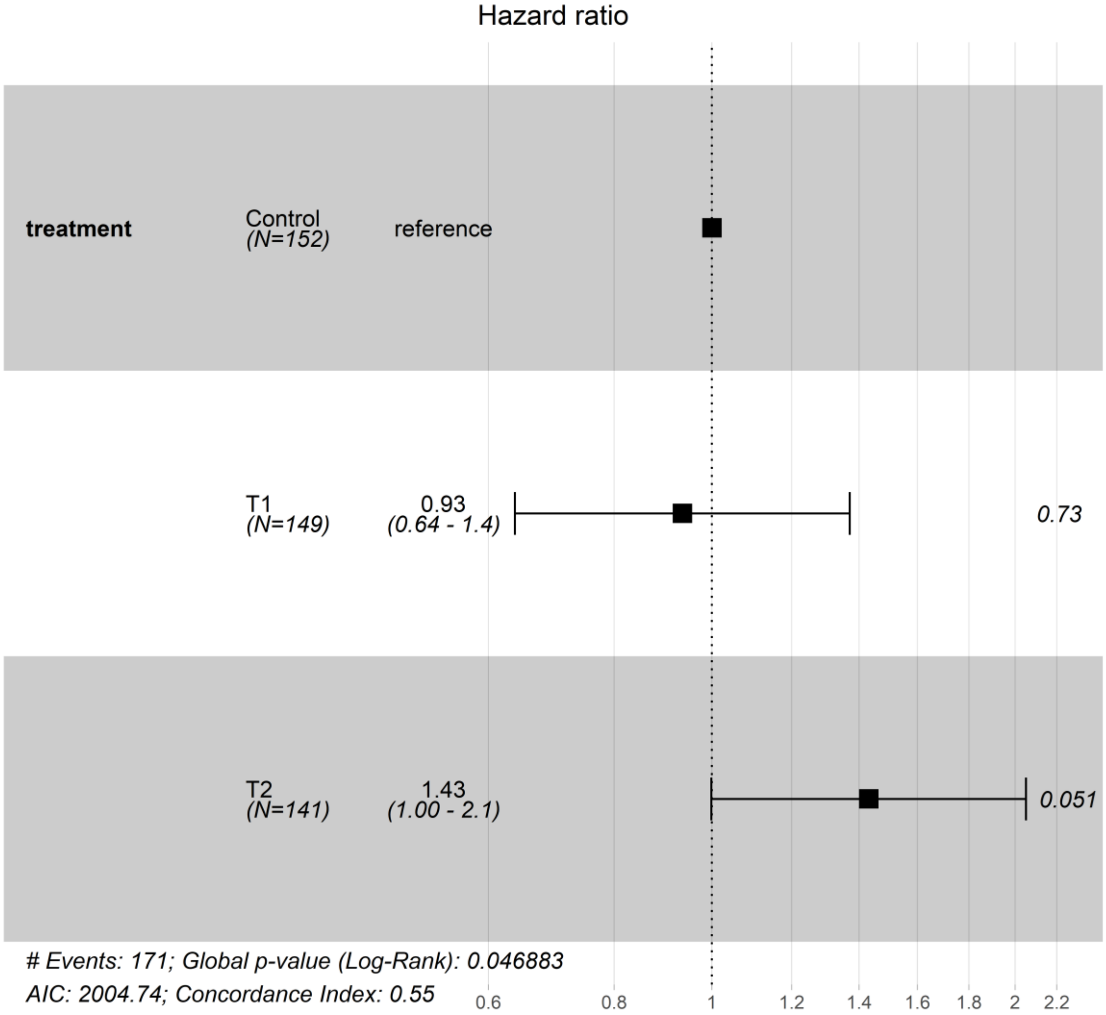


**Fig. S5** A Cox proportional hazards model was applied to determine the hazard ratio (HR) displayed as forest plot. Significant differences within those groups (treatment) were revealed close to the statistical threshold (global P = 0.047, log-rank test). A pairwise comparison, however, did not confirm these differences between the respective groups. With an HR of 0.93 for T1 and 1.43 for T2, the treated bees were not at risk of dying sooner when compared to the control (T1-C: P = 0.73, T2-C: P = 0.051, Log-rank test)

**Supplementary Figure S6:** Survival probability of untreated mini-hive replicates to justify pooling

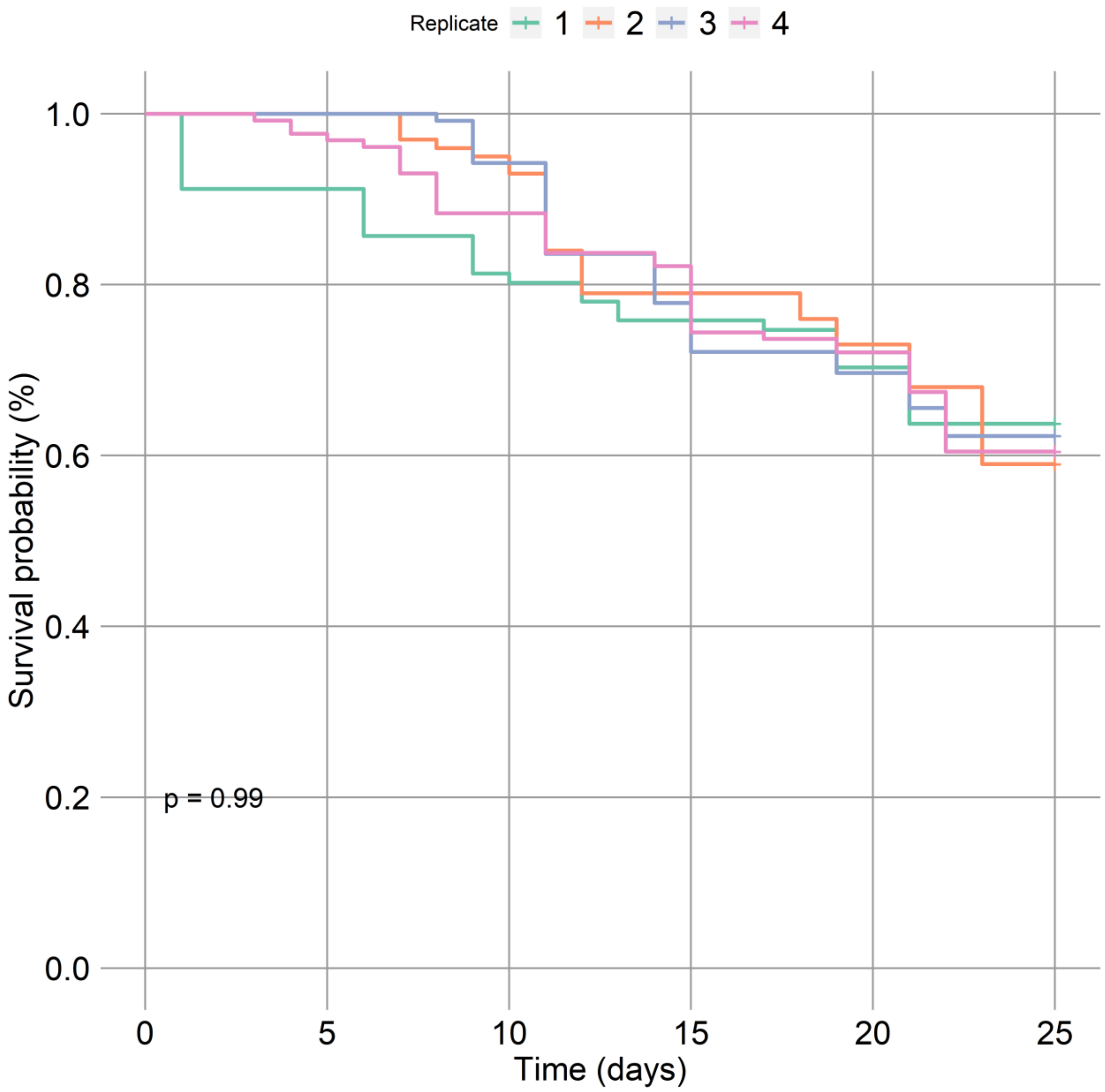


**Fig. S6** To justify pooling bees from the same groups but different mini-hives for survival analysis, these hives were evaluated separately treated as replicates. The test showed no significant differences ($P > 0.05$, Log-rank test).

### **Supplementary Method M S1:** Brood development and photographic assessment adopted from OECD 2007 and based on Schur et al. 2003

#### Brood termination rate

Based on the brood termination-rate the failure of individual eggs or larvae to develop is quantitatively assessed. For the calculation of the brood termination rate the observed cells are split into two categories:

- The bee brood in the observed cell reached the expected brood stage at the different assessment days or was found empty or containing an egg after hatch of the adult bee on BFD +22 → successful development
- The bee brood in the observed cell did not reach the expected brood stage at one of the assessment days or food was stored in the cell during BFD +5 to +16 → termination of the bee brood development

For the final calculation the number of cells, where a termination of the bee brood development was recorded, is summed up for each treatment and colony, is multiplied by 100 and divided by the number of cells observed in order to obtain of the brood termination rate in %.

#### Brood index

The brood index is an indicator of the bee brood development and facilitates a comparison between different treatments. The brood-index is calculated for each assessment day and colony. Therefore, the brood development in each cell will be checked starting from BFD 0 up to BFD +22. The cells are classified from 1 to 5 as described in paragraph 33, if the cells contain the expected brood stage at the different assessment days. If a cell does not contain the expected brood stage or food is stored in the cell during BFD +5 to +16 (see Table 4, OECD 2007) the cell has to be counted 0 (see Table 5, OECD 2007) at that assessment day and also on the following days, irrespective whether the cell is filled again with brood. This might require a further transformation of a value as described in paragraph 33. For the final calculation the values of all individual cells in each treatment, assessed at the same day, are summed up and divided by the number of observed cells in order to obtain the average brood index.

#### Compensation index

The compensation index is an indicator for recovery of the colony and will also be calculated for each assessment day and colony. The cells are classified from 1 to 5 as described in paragraph 33, solely based on the identified growth stage on the assessment days. By that, the compensation of bee brood losses will be included in the calculation of the indices. For the final calculation the values of all individual cells in each treatment, assessed at the same day, are summed up and divided by the number of observed cells in order to obtain the average compensation index.

**<u>Supplementary Method M S2:</u>** Analytical method and validation for glyphosate and AMPA

### Glyphosate analysis

### Preparation of feeding solution

A sample of 500 mg (approx. 400 µl) was weighed in a plastic tube (15 ml) and 9600 µl of the extracting agent (50 mM acetic acid/10 mM $Na_2EDTA$) were added. The tubes were closed and shaken thoroughly. Depending on the active substance concentration in the feeding solutions, these solutions were measured undiluted (control samples) or diluted to different extents. The dilutions were made with the extracting agent while adding the internal standards.

### Preparation of honey samples

A sample of approx. **1 g** was weighed in a plastic tube (15 ml) and a surrogate standard solution (20 µl Glufosinate (conc.: 2.5 ng/µl, corresponding to 10 pg/µl in the measuring solution)) and **4.3 ml** of the extracting agent (50 mM acetic acid/10 mM $Na_2EDTA$) were added to the sample. The tubes were closed and after homogenization using a Vortex-mixer left to stand for 30 minutes. Afterwards the tubes were shaken for one minute by hand, further 10 minutes with a horizontal shaker and then centrifuged for 5 minutes (1690 g). The entire supernatant was removed and filtered respectively cleaned using a Solid Phase Extraction (SPE) cartridge (OASIS HLB 6cc, 200 mg; Waters), to retain parts of the sugar in the sample. Before use, the SPE cartridge was conditioned with 2 ml methanol and 2 ml extracting agent and let run dry to not dilute the sample. 1000 µl of the sample extract were filled into a vial for measuring and 20 µl of an internal standard solution (glyphosate$^{13}C_2{}^{15}N$, AMPA$^{13}C^{15}N$, conc.: 1 ng/µl each, corresponding to 20 pg/µl in the measuring solution) were added. The measurements were started immediately after the completion of the extracts.

### Preparation of pollen samples

The samples were analyzed in the same way as described for honey with only one difference. **5.0 ml** of the extracting agent (50 mM acetic acid/10 mM $Na_2EDTA$) were added to the sample.

### Preparation of plant samples

A sample of **10 g** was weighed in a plastic cup (100 ml) and a surrogate standard solution (100 µL Glufosinate (conc.: 5 ng/µl, corresponding to 10 pg/µl in the measuring solution) and **50 ml** of the extracting agent (50 mM acetic acid/10 mM $Na_2EDTA$) were added. The cups were closed, left to stand for 30 minutes and then shaken for 20 minutes by means of a horizontal shaker.

Subsequently, the plant samples were crushed by means of a disperser and simultaneously extracted for 3 minutes. Then the samples were filtered using folded filters (particle retention 5 - 8 µm) into a 50 ml tube, or alternatively, the plant extracts were centrifuged (10 min, 1.690 g). **5 ml** of the filtered respectively centrifuged samples were transferred onto a conditioned (see preparation of honey samples) SPE-cartridge and filtered again respectively further cleaned. 1000 µl of the extract were prepared for measurements as described above (preparation of honey samples).

### Identification and quantification of the residues in the samples

LC-MS/MS was used for the identification and quantification of the target substances in the samples. Three multiple reaction monitoring (MRM) transitions were monitored for each analyte in order to confirm compound identity.

Reference standards in matrix and/or extracting agent were used for quantification, which was carried out according to the method of the internal standard. A large number of standards with the following concentrations were measured to create the calibration function: 0.1, 0.2, 0.5, 1, 5, 10, 25, 50, 100, 200, 500 and 1000 pg/µl). The results shown for the samples are averages of duplicate injections of sample extracts.

In undiluted and 1:10 diluted samples, the analyte contents were determined using matrix-matched standards. If samples had to be diluted 1:100 or 1:1000, the analytes were quantified using reference standards in extracting agent as matrix effects were sufficiently reduced by dilution.

The results for the surrogate standard glufosinate were used to control the analysis and were not included in the calculation of the analyte content in the samples.

### Equipment and measurement conditions

LC-MS/MS

The system used was a Nexera X2 HPLC system (SHIMADZU Corp., Kyoto, Japan) coupled to a triple quadrupole mass spectrometer Q TRAP 6500+ (SCIEX, Framingham, MA, USA) equipped with an electro spray ionization (ESI) source.

The mass spectrometric parameters were as follows:

- Scan type: sMRM
- Polarity: negative
- Ion spray voltage: - 4500 V
- Source temperature: 700 °C
- Curtain gas: 30 psi
- Nebulizer Gas (GS 1): 60 psi
- Heater Gas (GS 2): 60 psi
- Collision gas (nitrogen): high

The chromatographic separations were performed on an Acclaim Trinity Q1 column (3.0 x 100 mm; 3 µm, Thermo Fisher Scientific) with a pre-column SecurityGuard $C_{18}$ (3.0 x 4 mm, Phenomenex). The column oven temperature was set to 35 °C and the autosampler tray temperature was set to 15°C.

First, the samples were analyzed with the mobile phases (A) acetonitrile and (B) ultrapure water (0,055 µS/cm) with 50 mM ammonium formiate (adjusted to pH=2.9 with formic acid). The injection volume was 10 µl. The flow rates and gradient I and are shown in Tab. MS1.

Later the chromatographic conditions were optimized (Chamkasem & Vargo, 2017) and the samples analyzed with the mobile phases (A) ultrapure water and (B) ultrapure water with 50 mM ammonium formiate (adjusted to pH=2.9 with formic acid). A diverter valve between the LC column and the MS interface was used to direct the LC eluent to waste just before the AMPA peak (1.9 min) and after the glyphosate peak (3.5 min). The injection volume was 20 µl. The flow rates and gradient II are shown in Tab. MS2.

MRM transitions (negative mode) and compound dependent parameters are shown in Tab. MS3.

**Tab. MS1:** Gradient I

| Time (min) | Flow (ml/min) | Mobile phase A (%) | Mobile phase B (%) |
|---|---|---|---|
| 0.01 | 0.5 | 0 | 100 |
| 3.00 | 0.5 | 0 | 100 |
| 3.20 | 0.5 | 100 | 0 |
| 6.00 | 0.5 | 100 | 0 |
| 6.20 | 0.5 | 0 | 100 |
| 10.20 | 0.5 | 0 | 100 |

**Tab. MS2:** Gradient II

| Time (min) | Flow (ml/min) | Mobile phase A (%) | Mobile phase B (%) |
|---|---|---|---|
| 0.01 | 0.5 | 100 | 0 |
| 0.50 | 0.5 | 100 | 0 |
| 0.51 | 0.5 | 0 | 100 |
| 4.00 | 0.5 | 0 | 100 |
| 4.10 | 0.7 | 100 | 0 |
| 10.00 | 0.7 | 100 | 0 |

**Tab. MS3:** MRM-transitions (negative mode) and compound dependent parameters.

| Compound | Precursor Ion Q1 Mass (m/z) | Product Ion Q3 Mass (m/z) | DP (V) | CE (V) | CXP (V) |
|---|---|---|---|---|---|
| Glyphosate 1 | 168 | 63 | -30 | -26 | -7 |
| Glyphosate 2 | 168 | 150 | -30 | -14 | -9 |
| Glyphosate 3 | 168 | 81 | -30 | -20 | -9 |
| AMPA 1 | 110 | 63 | -15 | -24 | -7 |
| AMPA 2 | 110 | 81 | -15 | -18 | -9 |
| AMPA 3 | 110 | 79 | -15 | -34 | -9 |
| Glufosinate 1 (Surr) | 180 | 63 | -50 | -66 | -13 |
| Glufosinate 2 | 180 | 95 | -50 | -24 | -13 |
| Glufosinate 3 | 180 | 85 | -50 | -24 | -15 |

| Compound | Precursor Ion Q1 Mass (m/z) | Product Ion Q3 Mass (m/z) | DP (V) | CE (V) | CXP (V) |
|---|---|---|---|---|---|
| Glyphosate-$^{13}C_2{}^{15}N$ 1 | 171 | 63 | -50 | -26 | -7 |
| Glyphosate-$^{13}C_2{}^{15}N$ 2 | 171 | 153 | -50 | -14 | -9 |
| AMPA$^{13}C^{15}N$ 1 | 112 | 63 | -45 | -24 | -7 |
| AMPA$^{13}C^{15}N$ 3 | 112 | 79 | -45 | -38 | -9 |

DP =Declustering Potential, CE = Collision Energy, CXP = collision cell exit potential, Surr = surrogate standard

### Method validation - glyphosate

The validation study was performed to evaluate recoveries (REC), detection limits (LOD) and quantification limits (LOQ). Control samples of honey (used for honey stomach as well), pollen and plant material (phacelia) were fortified at different levels and 5 (4 respectively 7 in the case of plants) replicates extracted as described above. For the determination of recovery rates, detection and quantification limits, reference standards in solvent and matrix were prepared with the following concentration levels: 0.1, 0.25, 0.5, 1, 2.5, 5, 10, 25, 50, 100, 250 and 500 pg/µl.

The LOD was determined as the lowest concentration at which at least two MRM were detected, the peak signals of which were three times higher than the background noise of the chromatogram and the ratio of which was in the range of the required criteria (SANTE, 2019). The next highest concentration of the calibration standards above the detection limit was set as LOQ.

The results of the method validation procedure with honey, pollen and plant material are summarized in the following tables.

**Tab. MS4:** REC, RSD, LOD and LOQ of the target analytes in honey

| Honey | Fortification level | | | | | | | | | |
|---|---|---|---|---|---|---|---|---|---|---|
| | **50 µg/g** [1] (n=5) | | **250 µg/kg** (n=5) | | **25 µg/kg** (n=5) | | Standards in extracting agent | | Standards in honey matrix | |
| **Analytes** | REC [%] | RSD [%] | REC [%] | RSD [%] | REC [%] | RSD [%] | LOD [1] [µg/kg] | LOQ [1] [µg/kg] | LOD [µg/kg] | LOQ [µg/kg] |
| Glyphosate | 112 | 17 | 85 | 11 | 74 | 13 | 5.0 | 10 | 5.0 | 12.5 |
| AMPA | 110 | 14 | 85 | 9 | 77 | 10 | 2.5 | 5.0 | 2.5 | 5.0 |

[1] At a honey weight of 1 g and a 1:1000 dilution of the sample.

**Table MS5:** REC, RSD, LOD and LOQ of the target analytes in pollen

| **Pollen** | **Fortification level** | | | | | | | | | |
|---|---|---|---|---|---|---|---|---|---|---|
| | **250 µg/kg** (n=5) | | **50 µg/kg** (n=5) | | **25 µg/kg** (n=5) | | Standards in extracting agent | | Standards in pollen matrix | |
| **Analytes** | REC [%] | RSD [%] | REC [%] | RSD [%] | REC [%] | RSD [%] | LOD [µg/kg] | LOQ [µg/kg] | LOD [µg/kg] | LOQ [µg/kg] |
| Glyphosate | 79 | 13 | 80 | 4 | 148 | 18 | 5.0 | 10 | 12.5 | 25 |
| AMPA | 71 | 16 | 74 | 7 | 87 | 13 | 2.5 | 5.0 | 12.5 | 25 |

**Table MS6:** REC, RSD, LOD and LOQ of the target analytes in plant material

| **Plants (Phacelia)** | **Fortification level** | | | | | | | | | |
|---|---|---|---|---|---|---|---|---|---|---|
| | **250 µg/kg** (n=4) | | **50 µg/kg** (n=7) | | **25 µg/kg** (n=4) | | Standards in extracting agent | | Standards in phacelia matrix | |
| **Analytes** | REC [%] | RSD [%] | REC [%] | RSD [%] | REC [%] | RSD [%] | LOD [µg/kg] | LOQ [µg/kg] | LOD [µg/kg] | LOQ [µg/kg] |
| Glyphosate | 83 | 6 | 74 | 8 | 73 | 10 | 5.0 | 10 | 5.0 | 12.5 |
| AMPA | 83 | 5 | 77 | 8 | 82 | 9 | 2.5 | 5.0 | 12.5 | 25 |
